# *SMDT1* variants impair EMRE-mediated mitochondrial calcium uptake in patients with muscle involvement

**DOI:** 10.1101/2022.10.31.514480

**Authors:** Elianne P. Bulthuis, Merel J.W. Adjobo-Hermans, Bastiaan de Potter, Saskia Hoogstraten, Lisanne H.T. Wezendonk, Omar A.Z. Tutakhel, Liesbeth T. Wintjes, Bert van den Heuvel, Peter H.G.M. Willems, Erik-Jan Kamsteeg, M. Estela Rubio Gozalbo, Suzanne C.E.H. Sallevelt, Suzanne M. Koudijs, Joost Nicolai, Charlotte I. de Bie, Jessica E. Hoogendijk, Werner J.H. Koopman, Richard J. Rodenburg

**Affiliations:** Department of Biochemistry (286), Radboud Institute for Molecular Life Sciences, Radboud University Medical Centre, 6525 GA Nijmegen, The Netherlands; Human and Animal Physiology, Wageningen University & Research, 6700 AH Wageningen, The Netherlands; Translational Metabolic Laboratory, Department of Laboratory Medicine, Radboud University Medical Centre, 6525 GA Nijmegen, The Netherlands; Department of Human Genetics, Radboud University Medical Centre, 6525 GA Nijmegen, The Netherlands; Department of Pediatrics, Maastricht University Medical Centre, 6229 HX Maastricht, The Netherlands; Department of Clinical Genetics, Maastricht University Medical Centre, 6229 HX Maastricht, The Netherlands; Department of Neurology, Maastricht University Medical Centre, 6229 HX Maastricht, The Netherlands; Department of Genetics, University Medical Centre Utrecht, 3508 AB Utrecht, the Netherlands; Rudolf Magnus Institute of Neuroscience, University Medical Centre Utrecht, 3584 CG Utrecht, The Netherlands; Department of Pediatrics, Amalia Children’s Hospital, Radboud Center for Mitochondrial Medicine, Radboud Institute for Molecular Life Sciences, Radboud University Medical Center, 6525 GA Nijmegen, The Netherlands

**Author notes:** **Shared corresponding authors:** <u>Dr. W.J.H. Koopman</u>, Department of Pediatrics, Amalia Children’s Hospital, Radboud Institute for Mitochondrial Medicine (RCMM), Radboud Center for Molecular Life Sciences (RIMLS), Radboud University Medical Centre (Radboudumc), P.O. Box 9101, NL-6500 HB Nijmegen, The Netherlands. <u>Dr. R.J. Rodenburg</u>, Department of Pediatrics, Translational Metabolic Laboratory (774), Amalia Children’s Hospital, Radboud Institute for Mitochondrial Medicine (RCMM), Radboud Center for Molecular Life Sciences (RIMLS), Radboud University Medical Center (Radboudumc), P.O. Box 9101, NL-6500 HB Nijmegen, The Netherlands. Shared first authors (Elianne P. Bulthuis, Merel J.W. Adjobo-Hermans). **Author contributions:** clinical diagnosis, care and initiation of functional studies (MERG, SCEHS, SMK, JN, CIdB, JEH), whole exome sequencing (E-JK), Western blotting (BdP, EPB, LHTW, LTW), BN-PAGE analysis and generation of complemented cell lines (OAZT, LTW), calcium measurements (BdP, SH, EPB, MJWAH), immunofluorescence microscopy (BdP, EPB, MJWAH), data analysis (EPB, MJWAH, BdP, SH, WJHK), manuscript writing (EPB, MJWAH, BvdH, WJHK, RJR), overall supervision of the research (MJWAH, PHGMW, WJHK, RJR).

## Abstract

Ionic calcium (Ca^2+^) is a key messenger in signal transduction and its mitochondrial uptake plays an important role in cell physiology. This uptake is mediated by the mitochondrial Ca^2+^ uniporter (MCU), which is regulated by EMRE (essential MCU regulator) encoded by the *SMDT1* (single-pass membrane protein with aspartate rich tail 1) gene. This work presents the genetic, clinical and cellular characterization of two patients harbouring *SMDT1* variants and presenting with muscle problems. Analysis of patient fibroblasts and complementation experiments provide evidence that these variants lead to absence of EMRE protein, induce MCU subcomplex formation and impair mitochondrial Ca^2+^ uptake. However, the activity of the oxidative phosphorylation enzymes, mitochondrial morphology and membrane potential, as well as routine/ATP-linked respiration were not affected. We hypothesize that the muscle-related symptoms in the patients with *SMDT1* variants result from aberrant mitochondrial Ca^2+^ uptake.

## Introduction

Ionic calcium (Ca^2+^) is a second messenger that plays a central role in signal transduction, metabolism and cell survival (1)(**Berridge, 2016**). Under resting conditions, the free Ca^2+^ concentration in the cytosol ([Ca^2+^]_c_) is maintained at a low level (∼100 nM). In non-excitable cells like primary human skin fibroblasts (PHSFs), [Ca^2+^]_c_ is increased by Ca^2+^ release from the endoplasmic reticulum (ER) and/or influx of extracellular Ca^2+^ across the plasma membrane (PM) via Orai and transient receptor potential (TRP) channels (2)(**Parekh & Putney, 2005**). To allow signalling and prevent unwanted Ca^2+^-induced cytotoxicity, elevated [Ca^2+^]_c_ levels are returned to the resting level by ATPase action (3)(**Brini & Carafoli, 2009**). The latter actively transport Ca^2+^ back into the ER by action of sarco/endoplasmic reticulum Ca^2+^-ATPases (SERCAs) or remove it across the PM via PM-Ca^2+^-ATPases (PMCAs). In concert, activation of the above mechanisms can induce transient or oscillatory [Ca^2+^]_c_ changes (4)(**Dupont et al., 2011**). Mitochondria, classically recognized as important cellular ATP producers, play an important role in Ca^2+^ signalling since they can take up and release Ca^2+^ (5, 6)(**Nicholls, 2005; Valsecchi et al., 2009**).

Rapid mitochondrial Ca^2+^ uptake occurs at “mitochondrial synapses”, where mitochondria-ER tethering proteins bring the mitochondrial outer membrane (MOM) in close proximity to the ER membrane (7-10)(**Rutter & Rizzuto, 2000; De Brito & Scorrano, 2008; Picard, 2015; Csordás et al., 2018**). In addition, mitochondria can take up Ca^2+^ that enters the cell via PM-located Ca^2+^ channels (11)(**Montero et al., 2000**). Mitochondrial Ca^2+^ release occurs at a much slower rate than mitochondrial Ca^2+^ uptake, for instance by coupled Na^+^/Ca^2+^ (NCLX) and Na^+^/H^+^ (NHX) exchange (5, 12)(**Nicholls, 2005; Takeuchi et al., 2015**). In this way (**Supplementary Fig. 1**), mitochondrial Ca^2+^ uptake, release and buffering can directly alter the dynamics of free Ca^2+^ changes in the mitochondrial matrix ([Ca^2+^]_m_) and (locally) modulate the dynamics and signalling content of the [Ca^2+^]_c_ signal (5, 13, 14)(**Nicholls, 2005; Samanta et al., 2014; Mammucari et al., 2018**).

Inside the mitochondrial matrix compartment, Ca^2+^ can activate various enzymes of the tricarboxylic acid (TCA) cycle (15-18)(**Hansford & Chappell, 1967; Denton et al., 1972; Denton et al., 1978; McCormack & Denton, 1979**), leading to increased NADH and FADH_2_ production and stimulation of electron transport chain (ETC) activity. Moreover, Ca^2+^ might stimulate the generation of ATP by complex V (CV or F_o_F_1_-ATPase) of the mitochondrial oxidative phosphorylation (OXPHOS) system (19, 20)(**Das & Harris, 1990; Wescott et al., 2019**). Ca^2+^ was also demonstrated to (indirectly) activate the mitochondrial adenine nucleotide transporter (ANT), which exchanges ADP for ATP and is required for proper CV-mediated ATP generation and transfer of mitochondria-generated ATP to the cytoplasm (21)(**Mildaziene et al., 1995**). Alternatively, [Ca^2+^]_c_ can stimulate the rate of pyruvate-driven OXPHOS, mediated by the activity of the malate-aspartate shuttle (22)(**Szibor et al., 2020**). Irrespective of the mechanism, Ca^2+^-induced stimulation of mitochondrial function is considered an important pathway to match increased mitochondrial ATP production with increased cellular ATP demand during cell activation. Therefore, in addition to direct Ca^2+^ buffering, mitochondrial Ca^2+^ uptake and release can also modulate [Ca^2+^]_c_ kinetics by affecting the (local) ATP supply to SERCA and PMCA pumps (**Supplementary Fig. 1**).

The mitochondrial Ca^2+^ uniporter (MCU) complex is embedded in the mitochondrial inner membrane (MIM) and a prime mediator of mitochondrial Ca^2+^ uptake (23)(**Kamer & Mootha, 2015**). The MCU complex consists of the pore-forming mitochondrial Ca^2+^ uniporter (MCU) subunit, (a.k.a. MCUa; (24, 25)**Baughman et al., 2011; De Stefani et al., 2011**), the mitochondrial Ca^2+^ uptake 1 (MICU1) subunit (26)(**Perocchi et al., 2010**), the mitochondrial

Ca^2+^ uptake 2 (MICU2) subunit (27)(**Plovanich et al., 2013**) and the essential MCU regulator (EMRE) subunit (28)(**Sancak et al., 2013**). Depending on the experimental condition and tissue, evidence was provided that the MCU complex can additionally contain the dominant negative MCU paralog MCUb (28, 29)(**Raffaello et al., 2013; Sancak et al., 2013**), the MICU paralog MICU3 (27, 30)(**Plovanich et al., 2013; Paillard et al., 2017**), the mitochondrial calcium uniporter regulator 1 (MCUR1; (31)**Mallilankaraman et al., 2012a**) and the regulator SLC25A23 (32)(**Hoffman et al., 2014**). MICU1 and MICU2 inhibit mitochondrial Ca^2+^ uptake at low [Ca^2+^]_c_ and activate this uptake at higher [Ca^2+^]_c_ (33, 34)(**Pathak & Trebak, 2018; Mallilankaranaman et al., 2012b**). Cryo-EM analysis suggests that this phenomenon is mediated by [Ca^2+^]_c_-dependent blocking of the Ca^2+^-permeating pore by a single MICU1/MICU2 dimer (35, 36)(**Fan et al., 2020; Wang et al., 2020**). However, this hypothesis is challenged by other evidence suggesting that MICU subunits do not plug the pore, but positively modify its open probability at elevated cytosolic calcium concentrations (37)(**Garg et al., 2021**).

Dimerization of the MCU complex occurs between the amino (N)-terminal domains of two MCU subunits (35, 36)(**Fan et al., 2020; Wang et al., 2020**). In humans, the MCU complex is thought to consist of a tetrameric pore in which each MCU subunit binds a single EMRE subunit. However, *in vivo* evidence suggests that this 4:4 stoichiometry is not essential for gatekeeping and full channel activity (38)(**Payne et al., 2020**). Co-crystallization analysis of MCU and EMRE suggests that EMRE is required to keep the channel in an open conformation (39)(**Wang et al., 2019**). This is compatible with functional evidence in various knock-down/knock-out models (cells, flies and mice) identifying EMRE as a key regulator of the MCU complex and mitochondrial Ca^2+^ uptake (28, 39-43)(**Sancak et al., 2013; Tsai et al., 2016; Vais et al., 2016; Tufi et al., 2019; Wang et al., 2019; Liu et al., 2020**).

Although EMRE-negative flies exhibited reduced lifespan, their basal ATP levels and oxygen (O_2_) consumption were virtually unaffected (42)(**Tufi et al., 2019**). Likewise, whole-body and heart-specific O_2_ consumption were not affected and behavioural abnormalities were absent in *Smdt1*^*-/-*^ mice (43)(**Liu et al., 2020**). Remarkably, relative to wild type animals, *Smdt1*^*-/-*^ mice were capable of similar maximal work and responded normally to acute cardiac stress. Compatible with the *Smdt1*^*-/-*^ mouse model, mitochondrial Ca^2+^ uptake was also impaired in various whole-body and tissue-specific MCU knockout mouse models at different developmental stages (44-48)(**Pan et al., 2013; Holmström et al., 2015; Luongo et al., 2015; Kwong et al., 2015; Kwong et al., 2018**). These studies further demonstrated that whole-body *Mcu*^*-/-*^ mice possess a reduced exercise capacity (44)(**Pan et al., 2013**), but no impaired stress response in the heart (45)(**Holmström et al., 2015**). Skeletal muscle- and cardiomyocyte-specific inducible MCU knockout animals also displayed impaired responses to acute stress and reduced acute exercise performance (47, 48)(**Kwong et al., 2015; Kwong et al., 2018**). In comparison to *Smdt1*^*-/-*^ and *Mcu*^*-/-*^ models, whole-body knockout of *Micu1* induced a much more severe phenotype and (near) complete perinatal lethality in mice (49, 50)(**Antony et al., 2016; Liu et al., 2016**). In this model, surviving *Micu1*^*-/-*^ animals displayed an increased basal mitochondrial Ca^2+^ content, altered brain mitochondrial morphology and reduced cellular ATP levels (50)(**Liu et al., 2016**). Overall, whole-body knockout *Micu1*^*-/-*^ mice presented with combined neurological and muscular defects, compatible with human patients with *MICU1* mutations (51-53)(**Logan et al., 2014; Lewis-Smith et al., 2016; Musa et al., 2019**).

A previous large-scale whole exome sequencing (WES) study suggested a homozygous genetic variant in the EMRE-encoding *SMDT1* gene (NM_033318:exon2:c.255C>G:p.(Ser85Arg)) to be a potential pathogenic variant in an adult patient with alleged limb-girdle muscular dystrophy (LGMD) and dystonia (54)(**Monies et al., 2017**). However, in this study no cellular complementation or functional evidence was presented. Here we present the first integrated genetic, clinical and cellular characterization of two novel patients with *SMDT1* variants (P1-EMRE, P2-EMRE) and demonstrate in cells from patients with muscle problems that these *SMDT1*-gene defects induce absence of EMRE protein, formation of an MCU subcomplex and impairment of mitochondrial Ca^2+^ uptake.

## Results

### Patient genetic analysis

The MCU complex mediates mitochondrial Ca^2+^ uptake and contains four subunits (MCU, MICU1, MICU2 and EMRE). In humans, EMRE is encoded by the *SMDT1* gene. Using whole exome sequencing (WES) we here identified two patients carrying novel *SMDT1* variants (P1-EMRE, P2-EMRE; **Table 1**). The WES data was first filtered for potential pathogenic variants in all known disease genes. This did not result in any gene variant associated with disease phenotypes matching those of the two patients (data not shown). Next, the entire dataset was examined for candidate variants. Starting from the total number of variants detected, the number of variants that remained after each filter step is shown (**Supplementary Table 1**). Filtering was based on expected autosomal recessive inheritance and likely homozygosity of candidate variants. For P1-EMRE, this resulted in a single candidate variant in *SMDT1*. For P2-EMRE, this resulted in two candidate variants, of which one is located in the *PISD* (phosphatidylserine decarboxylase) gene (NM_002779.4(PSD):c.142C>T (p.(Arg48Trp)). Causality of the *PISD* variant is highly unlikely because the phenotype caused by known pathogenic variants in this gene, Liberfarb syndrome (OMIM 618889), does not match that of P2-EMRE. The other candidate variant is located in *SMDT1* (**Supplementary Table 1**). For comparison we included two control cell lines (CT1, CT2). Control CT2 also served as an experimental/technical quality control since it was extensively studied by us previously (*e*.*g*. (6, 55-59)**Visch et al., 2004; Visch et al., 2006a; Visch et al., 2006b; Valsecchi et al., 2009; Koopman et al., 2005; Distelmaier et al., 2009**). As a further control, we also included another patient (P3-MICU1), harbouring a known pathogenic variant (53)(**Musa et al., 2019**) in the *MICU1* gene (**Table 1**). This allowed further validation of our experimental approaches and results, since the impact of pathogenic *MICU1* variants is well studied (51, 52, 60)(**Logan et al., 2014; Lewis-Smith et al., 2016; Bhosale et al., 2017**).

**Table 1:**
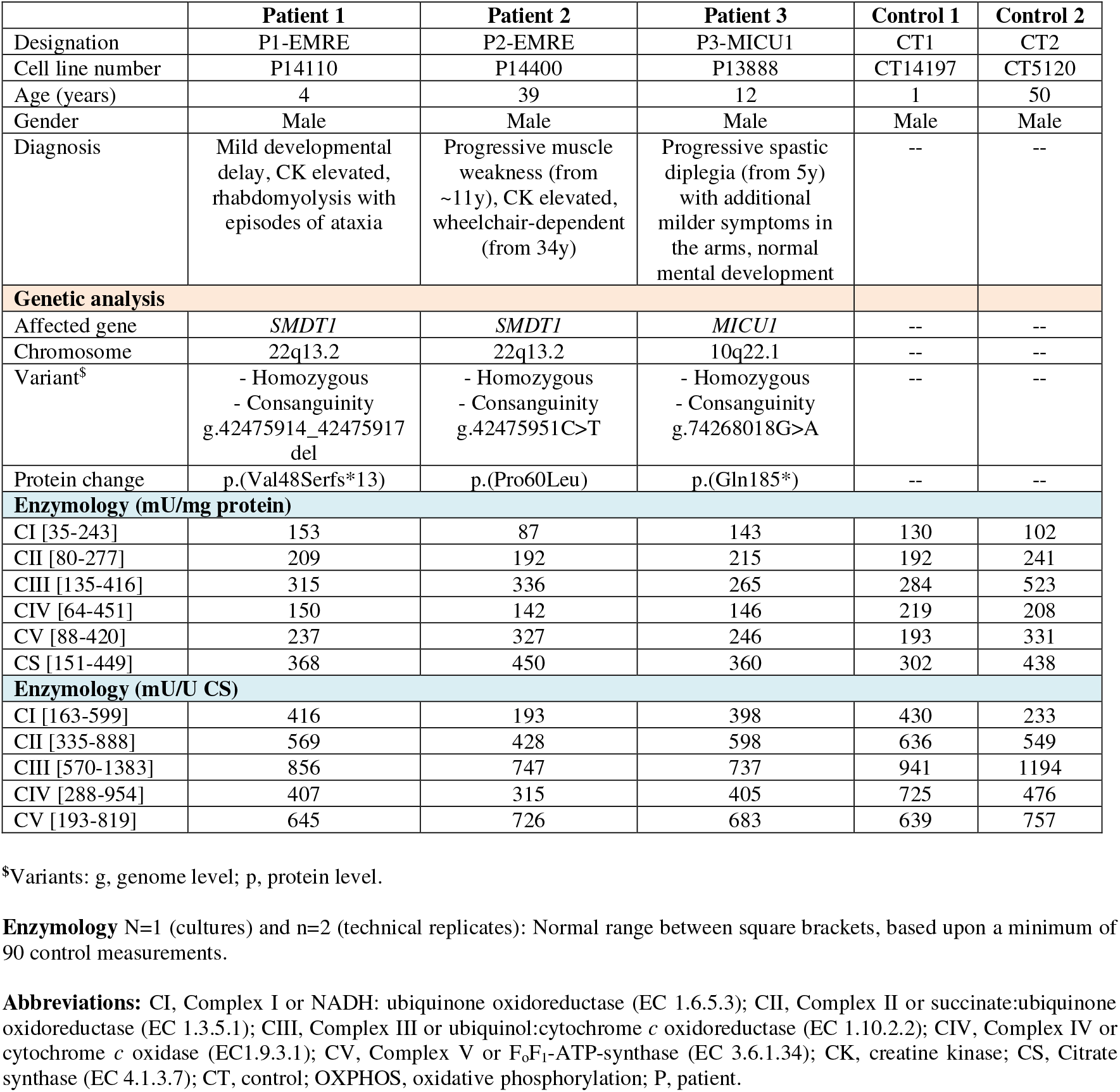
Cell lines, patient clinical phenotype, genetic analysis and enzymology.

|  | **Patient 1** | **Patient 2** | **Patient 3** | **Control 1** | **Control 2** |
| --- | --- | --- | --- | --- | --- |
| Designation | P1-EMRE | P2-EMRE | P3-MICU1 | CT1 | CT2 |
| Cell line number | P14110 | P14400 | P13888 | CT14197 | CT5120 |
| Age (years) | 4 | 39 | 12 | 1 | 50 |
| Gender | Male | Male | Male | Male | Male |
| Diagnosis | Mild developmental delay, CK elevated, rhabdomyolysis with episodes of ataxia | Progressive muscle weakness (from ~11y), CK elevated, wheelchair-dependent (from 34y) | Progressive spastic diplegia (from 5y) with additional milder symptoms in the arms, normal mental development | -- | -- |
| **Genetic analysis** |  |  |  |  |  |
| Affected gene | *SMDT1* | *SMDT1* | *MICU1* | -- | -- |
| Chromosome | 22q13.2 | 22q13.2 | 10q22.1 | -- | -- |
| Variant<sup>s</sup> | - Homozygous<br>- Consanguinity<br>g.42475914_42475917 del | - Homozygous<br>- Consanguinity<br>g.42475951C>T | - Homozygous<br>- Consanguinity<br>g.74268018G>A | -- | -- |
| Protein change | p.(Val48Serfs*13) | p.(Pro60Leu) | p.(Gln185*) | -- | -- |
| **Enzymology (mU/mg protein)** |  |  |  |  |  |
| CI [35-243] | 153 | 87 | 143 | 130 | 102 |
| CII [80-277] | 209 | 192 | 215 | 192 | 241 |
| CIII [135-416] | 315 | 336 | 265 | 284 | 523 |
| CIV [64-451] | 150 | 142 | 146 | 219 | 208 |
| CV [88-420] | 237 | 327 | 246 | 193 | 331 |
| CS [151-449] | 368 | 450 | 360 | 302 | 438 |
| **Enzymology (mU/U CS)** |  |  |  |  |  |
| CI [163-599] | 416 | 193 | 398 | 430 | 233 |
| CII [335-888] | 569 | 428 | 598 | 636 | 549 |
| CIII [570-1383] | 856 | 747 | 737 | 941 | 1194 |
| CIV [288-954] | 407 | 315 | 405 | 725 | 476 |
| CV [193-819] | 645 | 726 | 683 | 639 | 757 |

<sup>s</sup>Variants: g, genome level; p, protein level.
**Enzymology** N=1 (cultures) and n=2 (technical replicates): Normal range between square brackets, based upon a minimum of 90 control measurements.
**Abbreviations:** CI, Complex I or NADH: ubiquinone oxidoreductase (EC 1.6.5.3); CII, Complex II or succinate:ubiquinone oxidoreductase (EC 1.3.5.1); CIII, Complex III or ubiquinol:cytochrome *c* oxidoreductase (EC 1.10.2.2); CIV, Complex IV or cytochrome *c* oxidase (EC1.9.3.1); CV, Complex V or F<sub>0</sub>F<sub>1</sub>-ATP-synthase (EC 3.6.1.34); CK, creatine kinase; CS, Citrate synthase (EC 4.1.3.7); CT, control; OXPHOS, oxidative phosphorylation; P, patient.

### Patient clinical presentation

<u>Patient 1</u> (**P1-EMRE**) carries a homozygous 4bp deletion in the *SMDT1* gene (Chr22(GRCh37):g.42475914_42475917del; NM_033318.4:c.142_145del), leading to a frameshift within the first *SMDT1* exon and introducing a premature stop codon (p.(Val48Serfs*13)). This patient, a male, was born from healthy consanguineous parents (first cousins) at term with a normal birth weight. During pregnancy, no complications were observed and new-born screening revealed no abnormalities. The patient was admitted for the first time at the age of 13 months with an episode of acute ataxia. Cerebral MR imaging showed no abnormalities. However, serum creatine kinase (CK) level was increased (6176 U/l; highest reference value: 200 U/l). Both ataxia and CK levels normalized gradually. During the first years of life, the patient was admitted to the hospital several times because of episodes of intercurrent infections complicated by acute rhabdomyolysis (with elevated CK values). Dystonic posturing of the arms was observed during one of those admissions. Symptom-free intervals were characterized by highly variable but unambiguously elevated CK values (1597-12012 U/l). There are no known inherited diseases or rhabdomyolysis-related disorders in the family. An MRI performed at the age of 1 year was normal. At 4 years of age, the patient showed a mild developmental delay, partially explained by a language problem. The patient follows special education and has speech therapy. No visual or auditory problems have been noticed. Physical examination revealed no internal abnormalities. Normal muscle strength was observed, as well as normal symmetric tendon and plantar reflexes. Neither metabolite screening for metabolic diseases (amino acids, organic acids, acylcarnitines, purines and pyrimidines in blood and/or urine) nor cardiac screening (echocardiography) revealed any abnormalities. Lactate level was normal (1.3 mmol/l). Recently, at the age of 6 years he was admitted again because of recurrent periods of ataxia lasting for hours. These symptoms disappeared spontaneously. <u>Patient 2</u> (**P2-EMRE**) carries a homozygous single nucleotide substitution in the *SMDT1* gene (Chr22(GRCh37):g.42475951C>T; NM_033318.4:c.179C>T), changing a proline residue at position 60 into leucine (p.(Pro60Leu)). This patient, a currently 41 years old male, was born from healthy consanguineous parents (first cousins). He developed problems with running and climbing the stairs from the age of 11 years onwards. As of his twenties he also noticed weakness of the arms. Lactate level was normal (1.6 mmol/l). Muscle weakness was progressive over time and at age 34 years he became wheelchair-dependent. He reported a rapid deterioration of his symptoms after a gastroenteritis at age 30 years. Intellectual development was normal. There were no other family members with muscle disease. His past medical history revealed a bilateral orchidopexy (8 years) and testicular cancer (38 years). Neurological examination at age 36 years showed symmetrical winging scapulae, asymmetric wasting and weakness of all shoulder girdle muscle (MRC grade 4-, 4+) and finger extensors, and symmetrical weakness of all girdle and proximal muscles of the legs (0-4-) and foot dorsal flexors (grade 4). Over the years there was a gradual decrease of the vital capacity (60% of expected at age 36 years) without the need for non-invasive nocturnal ventilation up till now. Repeated cardiologic examinations were normal. Serum CK levels were elevated (456-3680 U/l). Muscle CT imaging (age 30 years) showed a predominant involvement of the subscapularis, long spinal muscles, thoracic wall muscles and thigh muscles. A muscle biopsy showed an increased fiber size variation, some endomysial thickening, necrotic and regenerating fibers, trabecular fibers, some enhanced subsarcolemmal staining in the Gomori but no evident ragged red fibers, no COX-negative fibers, best fitting a muscular dystrophy. Genetic testing over the years had shown normal findings for muscle disease multigene panels. SNP-array analysis was normal. Finally, WES was performed and identified a homozygous *SMDT1* variant. <u>Patient 3</u> (**P3-MICU1**) carries a homozygous single nucleotide substitution in the *MICU1* gene (Chr10(GRCh37):g.74268018G>A; NM_006077.3:c.553C>T), changing residue Q185 into a stop codon (p.(Gln185*)). The patient included in our study, a 12-year-old male, was born from healthy consanguineous parents. From the age of 5 years onwards, he displayed progressive spastic diplegia with additional milder symptoms in the arms. Serum CK levels were increased, whereas lactate, alanine and organic acid levels were normal. His mental development was normal. No information was available with respect to reflexes or the brain at the level of MRI.

### SMDT1 gene defects induce absence of EMRE and formation of an MCU subcomplex

Cellular analyses were performed by comparing primary human skin fibroblasts (PHSFs) of the patients (P1-EMRE, P2-EMRE, P3-MICU1) with two control cell lines from healthy donors (CT1, CT2). Enzymological analysis demonstrated that none of these cell lines displayed abnormalities in OXPHOS complex and CS activities (**Table 1**). SDS-PAGE and Western blot (WB) analysis (**Fig. 1A** and **Supplementary Fig. 2A-B**) revealed that the MCU protein was present in all cell lines (**Fig. 1A**; black boxes), whereas MICU1 was not detected in P3-MICU1 cells. In case of EMRE, a WB-optimized antibody was used revealing bands in mitochondrial fractions of CT1 and CT2 cells (**Fig. 1A**; green boxes). Mitochondrial EMRE protein was not detected in P1-EMRE, P2-EMRE and P3-MICU1 cells (**Fig. 1A**; red boxes). Blue Native (BN)-PAGE analysis of mitochondria-enriched fractions with an MCU-specific antibody (**Fig. 1B, Supplementary Fig. 2C** and **Supplementary Fig. 5B**) demonstrated the presence of an MCU subcomplex of lower MW in all three patient cell lines. Although no mitochondrial marker protein (VDAC1) was detected in the cytosolic fractions, the latter did exhibit MICU1-positive bands for all cell lines except P3-MICU1 (**Fig. 1A**). Cytosolic MICU1 signals were highest for P1-EMRE and P2-EMRE, suggesting that absence of EMRE destabilizes the MCU complex (supported by the BN-PAGE data in **Fig. 1B**), and that this destabilization induces translocation of MICU1 to the cytosol. To better understand the subcellular localization of EMRE, immunofluorescence (IF) microscopy was performed using an IF-optimized EMRE antibody (**Fig. 2; Supplementary Fig. 3B-C**). This revealed a punctate mitochondrial EMRE staining in CT1 and CT2 cells. A similarly patterned but apparently lower EMRE signal was observed in P3-MICU1 cells (**Supplementary Fig. 3D**). In contrast, no mitochondrial EMRE signal was detected in P1-EMRE and P2-EMRE cells (**Fig. 2; Supplementary Fig. 3C**). Particularly in cells displaying no (P1-EMRE, P2-EMRE) or low (P3-MICU1) mitochondrial EMRE staining a strong nuclear fluorescence was detected (**Fig. 2; Supplementary Fig. 3B-D**). This is likely due to non-specific binding of the primary EMRE antibody, since such a staining pattern was absent when only the secondary antibody was used (**Supplementary Fig. 3A**). Taken together, these results suggest that the *SMDT1* gene defects induce absence of EMRE but not of MCU and MICU1 in P1-EMRE and P2-EMRE cells. In contrast, the *MICU1* defect is associated with absence of MICU1 protein and lower levels of EMRE.

**Figure 1:**
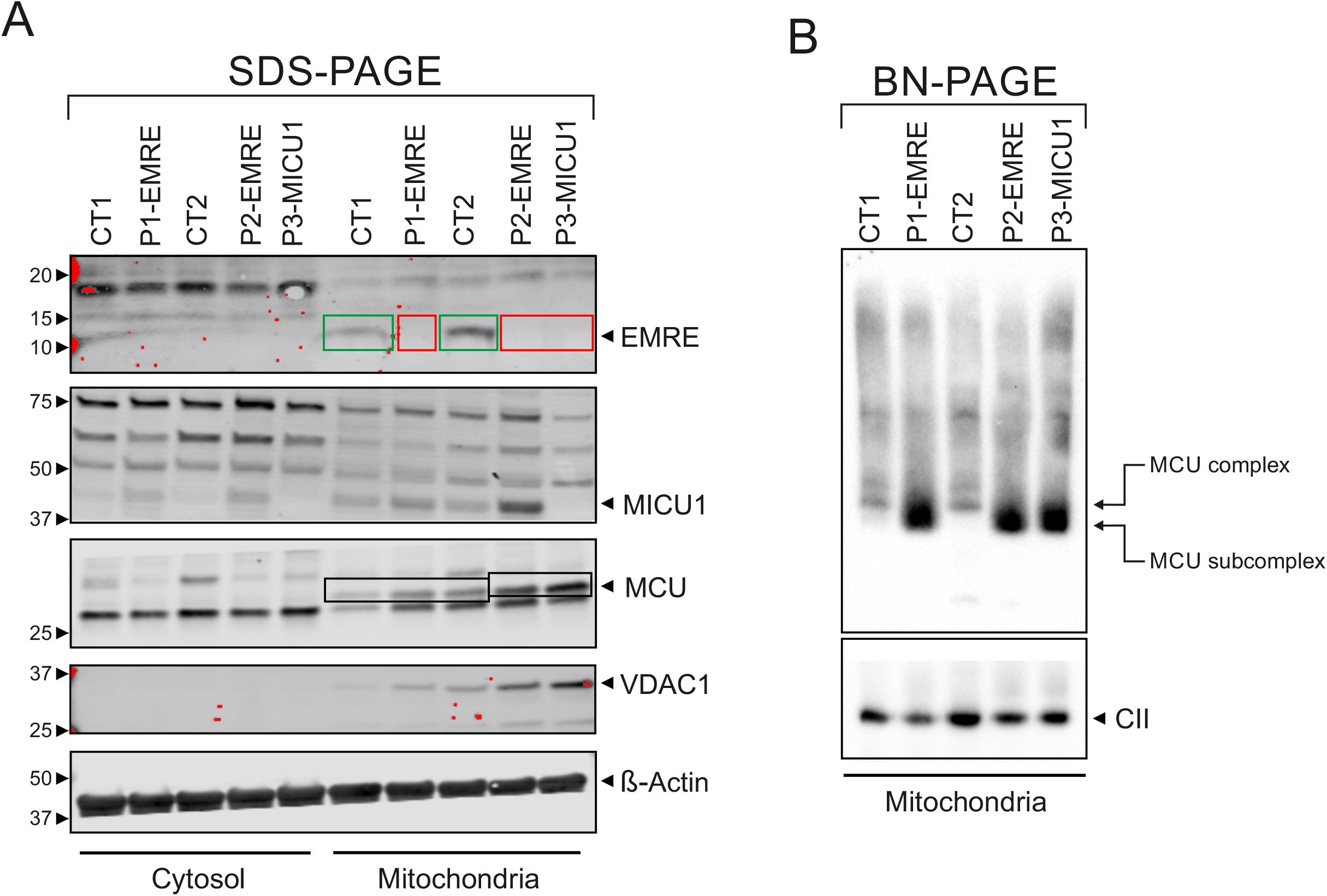
*SMDT1* defects hamper formation of the MCU complex in patient fibroblasts. (**A**) SDS-PAGE and Western blot analysis of cytosolic and mitochondrial fractions isolated from PHSFs from control subjects (CT1, CT2) and the three patients (P1-EMRE, P2-EMRE, P3-MICU1). Antibodies against EMRE, MICU1, MCU, VDAC1 (mitochondrial marker) and β-actin (cytosolic marker) were used. Two EMRE-positive bands were detected in CT1 and CT2 cells (green boxes), whereas such bands were not detected in P1-EMRE, P2-EMRE and P3-MICU1 cells (red boxes). The saturated (red) pixels on the left of the blot are derived from the MW markers (also see **Supplementary figure 2**). Numerals indicate molecular weight in kDa. (**B**) Native gel (BN-PAGE) and Western blot analysis of mitochondria-enriched fractions isolated from PHSFs and stained with antibodies against MCU and OXPHOS complex II (CII; loading control). The fully assembled MCU complex (“MCU complex”) and the subcomplex lacking the EMRE or MICU1 subunit (“MCU subcomplex”) are indicated.

**Figure 2:**
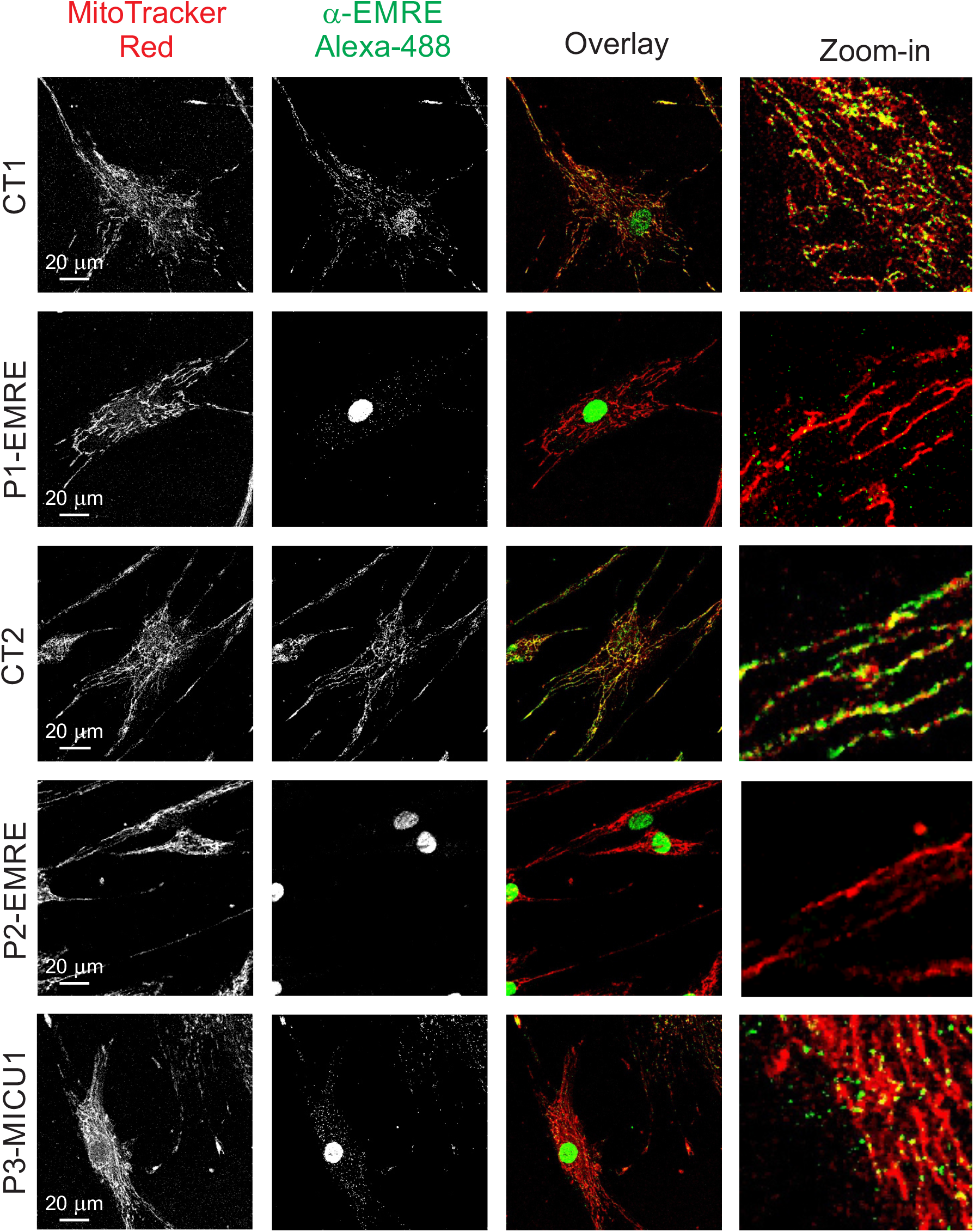
*SMDT1* defects trigger loss of the EMRE protein in patient fibroblasts. Typical confocal microscopy images of control (CT1, 19 cells; CT2, 8 cells) and patient-derived PHSFs (P1-EMRE, 7 cells; P2-EMRE, 43 cells; P3-MICU1, 14 cells) co-stained with the mitochondrial marker MitoTracker Red (red) and anti-EMRE/Alexa-488 antibodies (green). Images were processed for visualization purposes by subsequent application of a linear contrast stretch (LCS) operation, a median filter (3×3; single pass) and a second LCS operation.

**Figure 3:**
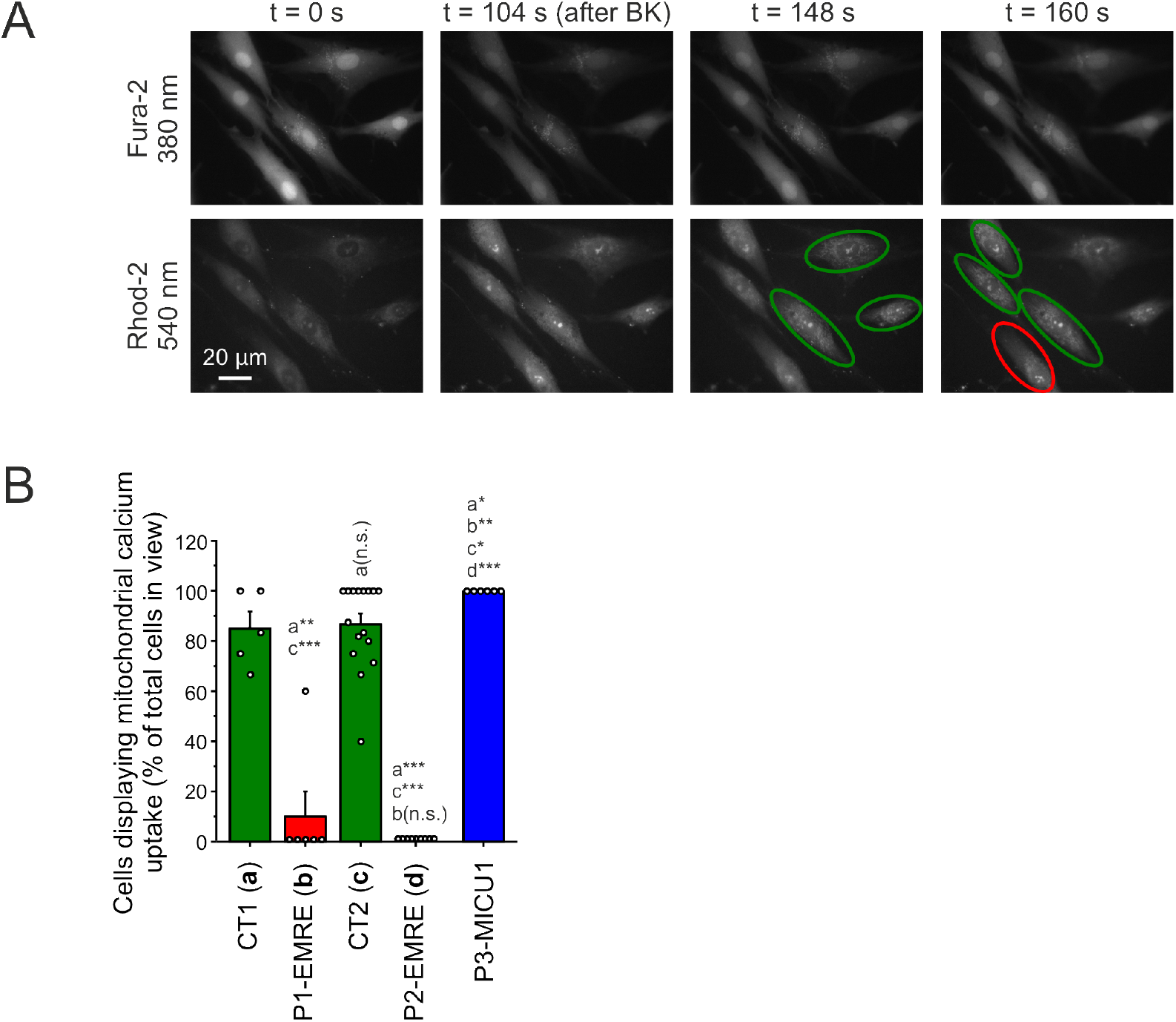
*SMDT1* defects prevent mitochondrial Ca^2+^ uptake and apparently reduce the amplitude of cytosolic *Ca*^*2+*^ signals in hormone-stimulated patient fibroblasts. (**A**) Bradykinin (BK)-induced increases in cytosolic Ca^2+^ concentration ([Ca^2+^]_c_) and mitochondrial Ca^2+^ concentration ([Ca^2+^]_m_) were detected in PHSFs co-loaded with fura-2 ([Ca^2+^]_c_) and rhod-2 ([Ca^2+^]_m_). This typical example illustrates this strategy for CT1 cells. At the start of the recording (t = 0 s), the fura-2 fluorescence signal (380 nm excitation) is high and the rhod-2 fluorescence signal is low, indicating a low [Ca^2+^]_c_ and [Ca^2+^]_m_, respectively. Upon BK addition (t = 104 s) the fura-2 signal decreased and the nuclear/cytosolic rhod-2 signal increased, demonstrating an increase in [Ca^2+^]_c_. Cells that did not show a [Ca^2+^]_c_ increase (*i*.*e*. did not respond to BK) were omitted from further analysis. Following BK addition, the fura-2 signal gradually increased (t = 148 s and t = 160 s), compatible with a gradual decrease in [Ca^2+^]_c_. During this decrease mitochondrial rhod-2 fluorescence became clearly visible, indicating that [Ca^2+^]_m_ increased. To be certain that no mitochondrial rhod-2 signals were missed due to the small mitochondrial size, the focus was continuously adjusted manually during the measurement. To analyse the experiment, cells were manually scored for positive mitochondrial Ca^2+^ uptake when a mitochondrial rhod-2 signal was observed in one of the axial images (*e*.*g*. green ovals at t = 148 s and t = 160 s). In this example, one cell was scored negative for mitochondrial Ca^2+^ uptake (red oval at t = 160 s). Hence, 5 out of 6 cells (83.33 %) in the field of view (FOV) were scored positive in this example. (**B**) Analysis of BK-stimulated mitochondrial Ca^2+^ uptake in fura-2/rhod-2 co-loaded cells, as explained in panel A. Each symbol represents a single FOV. This data was obtained in at least N=2 independent experiments for n=26 cells (CT1; 5 dishes/FOVs), n=89 (CT2; 16 dishes/FOVs), n=39 (P1-EMRE; 6 dishes/FOVs), n=39 (P2-EMRE; 9 dishes/FOVs) and n=35 (P3-MICU1; 6 dishes/FOVs).

### SMDT1 gene defects impair mitochondrial Ca^2+^ uptake in hormone-stimulated cells

Functionally, EMRE plays a central role in MCU-mediated mitochondrial Ca^2+^ uptake. Hence, we applied live-cell fluorescence microscopy to determine whether *SMDT1* gene defects affected mitochondrial Ca^2+^ uptake. To this end, PHSFs were co-stained with the fluorescent reporter molecules fura-2 and rhod-2 (**Fig. 3A**). In these cells, fura-2 resides in the cytosol and nucleoplasm, whereas rhod-2 is predominantly localized in the mitochondrial matrix but also is present at a low concentration in the cytosol/nucleoplasm (57)(**Visch et al., 2006b**). In this way the fura-2 signal can be used to detect changes in [Ca^2+^]_c_, whereas the rhod-2 signal reports on changes in both [Ca^2+^]_c_ (by its nuclear/cytosolic fluorescence signal) and [Ca^2+^]_m_ (mitochondrial fluorescence). Mitochondrial Ca^2+^ uptake in PHSFs is stimulated by acute extracellular application of the hormone Bradykinin (BK; 1 μM; (6, 55)**Visch et al., 2004; Valsecchi et al., 2009**). This involves Ca^2+^ release from the endoplasmic reticulum (ER) via the G-protein coupled receptor (GPCR) and inositol 1,4,5-trisphosphate (InsP_3_)-mediated pathway (**Supplementary Fig. 1**). In our experience, BK stimulation not necessarily induces a [Ca^2+^]_c_ increase in all PHSFs. This necessitates the simultaneous monitoring of the nuclear fluorescence signals of the Ca^2+^-free form of fura-2 and rhod-2, as well as the mitochondrial rhod-2 signal (**Fig. 3A**). In addition, because mitochondrial rhod-2 signals are absent in PHSFs prior to BK stimulation (57, 61)(**Visch et al., 2006b; Koopman et al., 2008**), mitochondria are not visible at the start of the experiment (**Fig. 3A**; t = 0 s). Therefore, we continuously refocused during image acquisition to allow proper detection of BK-induced increases in mitochondrial rhod-2 signal. Although this approach precludes kinetic analysis of the fluorescence signals, it allowed proper scoring of cells displaying mitochondrial Ca^2+^ uptake (**Fig. 3A, Supplementary Movie 1-2**). This demonstrated that all cells were BK-responsive (CT1: 36 cells, CT2: 99 cells, P1-EMRE: 39 cells, P2-EMRE: 39 cells, P3-MICU1: 35 cells). For unknown reasons some control cells exhibited a mitochondrial rhod-2 fluorescence signal prior to BK stimulation and therefore were excluded from further analysis (CT1: 10 cells; CT2: 10 cells). Rhod-2 analysis by blinded scoring demonstrated that the fraction of cells in a given field of view (FOV) displaying BK-induced mitochondrial Ca^2+^ uptake was high in CT1, CT2 and P3-MICU1 cells, whereas this fraction was virtually zero in P1-EMRE and P2-EMRE cells (**Fig. 3B, Supplementary Movie 1-2**). Independent blind scoring by another researcher yielded similar results (**Supplementary Fig. 4A**). Of note, the majority of P3-MICU1 cells (25 out of 35; 71%) already displayed an increased mitochondrial rhod-2 signal in the absence of BK stimulation. These results demonstrate that BK-stimulated mitochondrial Ca^2+^ uptake is impaired in P1-EMRE and P2-EMRE cells.

### Complementation with the wild type SMDT1 gene rescues MCU subcomplex formation and impaired mitochondrial Ca^2+^ uptake

To better causally link absence of EMRE (**Fig. 1A**) to MCU subcomplex formation (**Fig. 1B, Supplementary Fig. 2C** and **Supplementary Fig. 5B**) and impaired mitochondrial Ca^2+^ uptake (**Fig. 3B**), P1-EMRE cells were complemented with the wild type *SMDT1* gene. We selected P1-EMRE cells for these rescue experiments since these harboured an *SMDT1* premature stop codon and therefore were unable to synthesize full-length EMRE protein. Genetic labelling of the EMRE protein with a V5 tag followed by SDS-PAGE and WB analysis demonstrated that the wild type EMRE protein was expressed in the cells (**Fig. 4A** and **Supplementary Fig. 5A**). Microscopy analysis with the IF-optimized EMRE antibody revealed a punctate mitochondrial staining in ∼30% of the P1-EMRE+SMDT1-V5 cell population (**Fig. 4B**). The latter was accompanied by partial restoration of the fully assembled MCU complex (**Fig. 4C** and **Supplementary Fig. 5B**) and partial restoration of BK-stimulated mitochondrial Ca^2+^ uptake (**Fig. 4D**). The latter was confirmed by independent blind scoring by another researcher (**Supplementary Fig. 4B**). As expected, a similar rescue was not observed in P1-EMRE transduced with GFP-V5 (**Fig. 4D**). These results strongly suggest that the partial restoration of the MCU complex (**Fig. 4C**) and mitochondrial Ca^2+^ uptake (**Fig. 4D**) is due to only a subfraction of the P1-EMRE cells expressing the SMDT1-V5 construct (**Fig. 4B**). We conclude that absence of EMRE is responsible for MCU subcomplex formation and impaired mitochondrial Ca^2+^ uptake in P1-EMRE cells.

**Figure 4:**
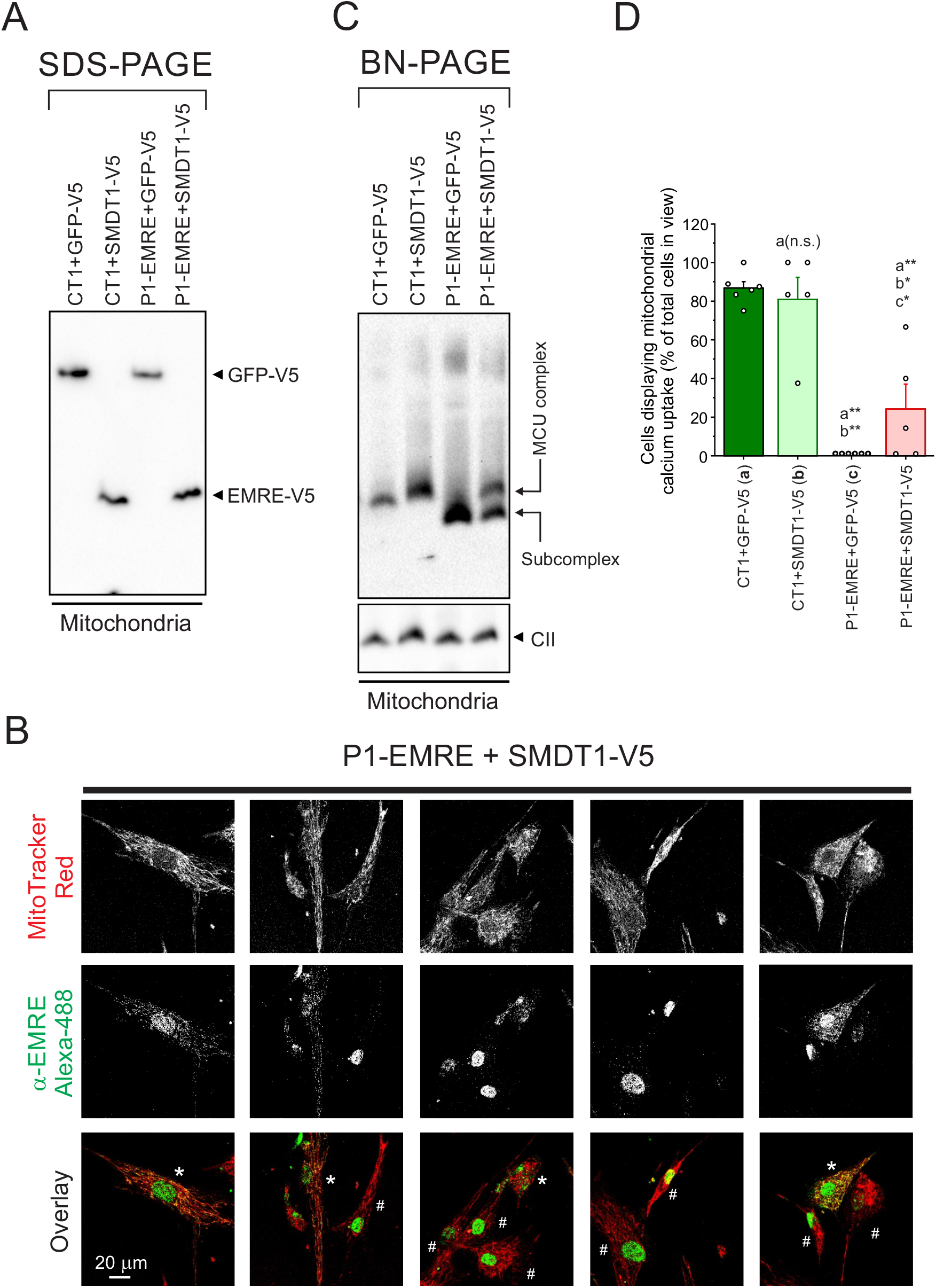
Rescue of the *SMDT1* defect partially restores formation of the MCU complex and hormone-induced mitochondrial Ca^2+^ uptake in patient fibroblasts. (**A**) SDS-PAGE and Western blot analysis of mitochondria-enriched fractions of CT1 and P1-EMRE cells lentivirally transduced with V5-tagged green fluorescent protein (GFP-V5) or the non-mutated *SMDT1* gene (+SMDT1-V5). An antibody directed against the V5 epitope was used for detection. (**B**) Typical confocal microscopy images of P1-EMRE cells transfected with SMDT1-V5 (In total 40 cells were visually inspected). Cells were co-stained with the mitochondrial marker MitoTracker Red (red) and anti-EMRE/Alexa-488 antibodies (green). Images were processed for visualization purposes by subsequent application of a linear contrast stretch (LCS) operation, a median filter (3×3; single pass) and a second LCS operation. SMDT1-positive and negative cells are marked by * and #, respectively. (**C**) Native gel (BN-PAGE) and Western blot analysis of mitochondria-enriched fractions for the conditions in panel A using antibodies against MCU and OXPHOS complex II (CII; loading control). The fully assembled MCU complex (“MCU complex”) and an MCU subcomplex (“MCU subcomplex”) are indicated. (**D**) Analysis of BK-stimulated mitochondrial Ca^2+^ uptake in fura-2/rhod-2 co-loaded cells. Each symbol represents a single field of view (FOV). This data was obtained in N=2 independent experiments for n=48 responding out of 48 cells (CT1+GFP-V5; 6 dishes/FOVs), n=35 out of 43 (CT1+SMDT1-V5; 5 dishes/FOVs), n=40 out of 40 (P1-EMRE+GFP-V5; 6 dishes/FOVs) and n=41 out of 47 (P1-EMRE+SMDT1-V5; 5 dishes/FOVs). **Statistics:** in panel D significant differences with the indicated columns (a,b,c) are marked by **(p<0.01) and *(p<0.05). N.s. indicates not significant.

### SMDT1 genetic variants do not alter mitochondrial membrane potential

Mitochondrial Ca^2+^ uptake is highly dependent on the trans-MIM membrane potential (ΔΨ; (62, 63)**Gunter & Gunter, 1994; De Stefani et al., 2016**). Therefore, the impaired mitochondrial Ca^2+^ uptake in P1-EMRE and P2-EMRE cells might be affected by the *SMDT1* genetic variants inducing (partial) ΔΨ depolarisation prior to BK addition, thereby decreasing the driving force for mitochondrial Ca^2+^ entry. To investigate this possibility, we assessed the mitochondrial accumulation of the fluorescent cation TMRM as a semi-quantitative readout of ΔΨ in CT1, CT2, P1-EMRE and P3-MICU1 cells using a previously described protocol (61, 64, 65)(**Koopman et al., 2008; Distelmaier et al., 2008; Esteras et al., 2020**). Mitochondrial TMRM fluorescence did not differ between CT1 and P1-EMRE cells, whereas P3-MICU1 displayed a higher TMRM signal than CT1 and P1-EMRE (**Fig. 5A**). This argues against ΔΨ depolarisation being responsible for impaired mitochondrial Ca^2+^ uptake in *SMDT1*-patient cells.

**Figure 5:**
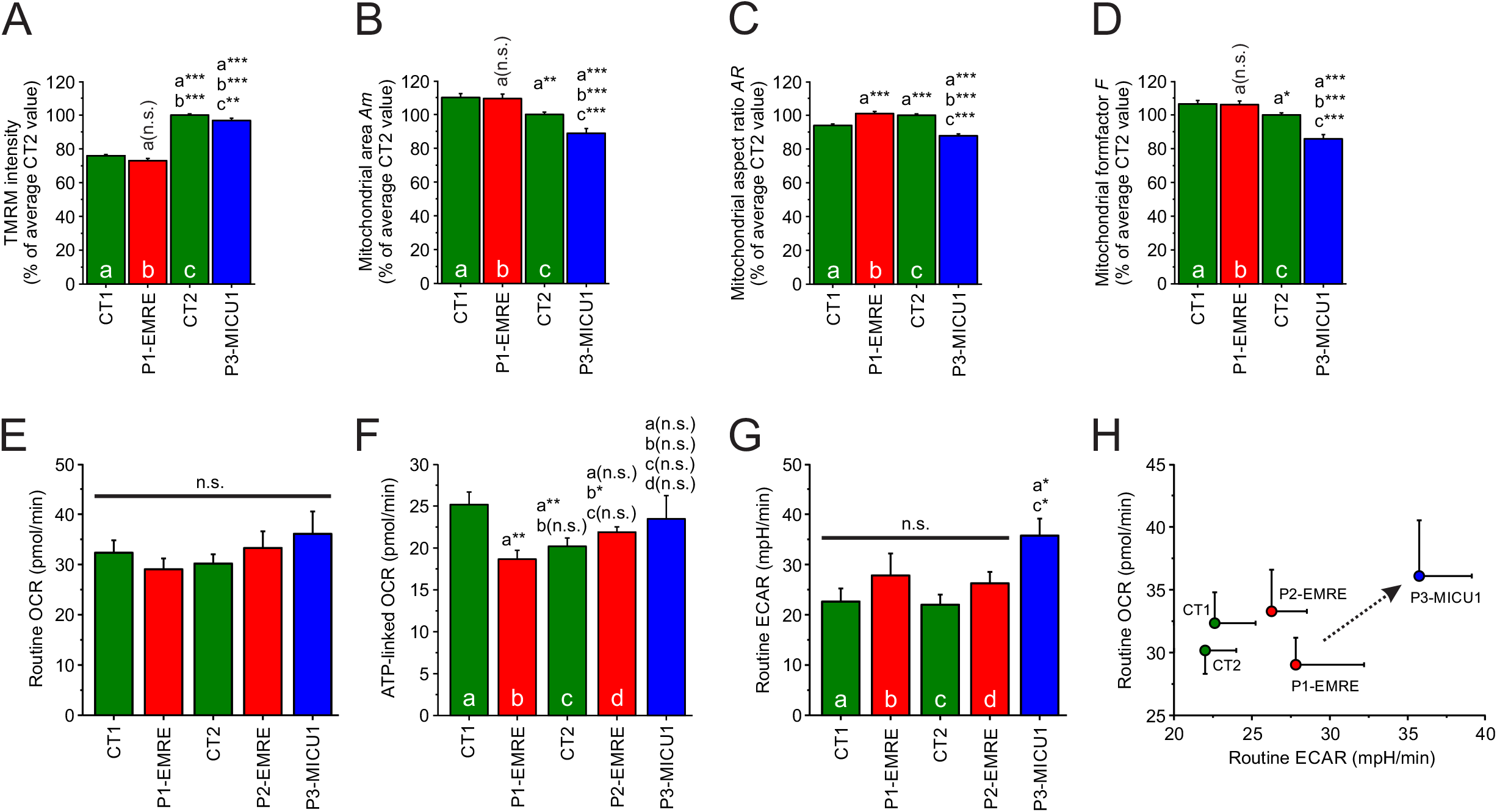
*SMDT1* defects do not affect mitochondrial membrane potential, mitochondrial morphology and oxygen consumption rate in primary patient fibroblasts. (**A**) Mitochondrial TMRM fluorescence intensity (expressed as percentage of the average CT2 signal measured on the same day) in CT1, CT2, P1-EMRE and P3-MICU1 cells. (**B**) Size of TMRM-positive objects (*Am*; expressed as percentage of the average CT2 value measured on the same day). (**C**) Same as panel B, but now for mitochondrial aspect ratio (*AR*). (**D**) Same as panel B, but now for mitochondrial formfactor (*F*). (**E**) Routine (basal) oxygen consumption rate (OCR) in CT1, CT2, P1-EMRE, P2-EMRE and P3-MICU1 cells. (**F**) ATP-linked OCR. (**G**) Routine extracellular acidification rate (ECAR). (**H**) Relationship between the OCR and ECAR data in panel E and G. The arrow suggests that P3-MICU1 cells are metabolically more active (*i*.*e*. display an increased OCR and ECAR). **Statistics:** For panel A,B,C,D the number of experiments (N) and replicates (n) equalled: CT1 (N>3, n=284), CT2 (N>3, n=414), P1-EMRE (N>3, n=162), P3-MICU1 (N>3, n=161). For panel E,F,G,H these numbers equalled: CT1 (N=3, n=16), CT2 (N=5, n=30), P1-EMRE (N=3, n=17), P2-EMRE (N=2, n=10), P3-MICU1 (N=3, n=18). Significant differences with the indicated columns (a,b,c,d) are marked by ***(p<0.001), **(p<0.01) and *(p<0.05). N.s. indicates not significant.

### SMDT1 genetic variants do not alter mitochondrial morphology

Alterations in mitochondrial morphology have been demonstrated to influence ER-to-mitochondria Ca^2+^ transfer (8, 66)(**Szabadkai et al., 2004; de Brito & Scorrano, 2008**) and it was previously demonstrated that absence of MICU1 induces a fragmented mitochondrial phenotype (51, 67)(**Logan et al., 2014; Gottschalk et al., 2019**). To determine if absence of EMRE altered mitochondrial morphology as well, we analysed TMRM-stained cells using a previously described approach (61)(**Koopman et al., 2008**). This delivered information on the area of individual mitochondrial objects (*Am*; a measure of mitochondrial size), mitochondrial aspect ratio (*AR*; a measure of mitochondrial length) and mitochondrial formfactor (*F*; a combined measure of mitochondrial length and degree of branching). P3-MICU1 cells displayed lower *Am, AR* and *F* values than control and P1-EMRE cells (**Fig. 5B-C-D**), indicating that mitochondria are smaller and more circular in these cells. In contrast, P1-EMRE cells displayed no major differences relative to control cells, except for a slight increase in *AR*, as compared to CT1 (**Fig. 5C**). This demonstrates that the *MICU1* gene defect induces a fragmented mitochondrial phenotype, whereas the *SMDT1* mutation does not alter mitochondrial morphology.

### SMDT1 genetic variants do not alter cellular oxygen consumption rate and extracellular acidification rate

Finally, we analysed if mitochondrial function was affected in control and patient cells. In these experiments, the routine (resting) cellular oxygen consumption rate (OCR; **Fig. 5E**) and extracellular acidification rate (ECAR; **Fig. 5G**) were used as a readout of OXPHOS and glycolytic activity, respectively (68)(**Divakaruni et al., 2014**). Routine OCR and ECAR did not differ between any of the cell lines, with the exception of P3-MICU1, which displayed an increased ECAR. With respect to ATP-linked OCR (**Fig. 5F**), P1-EMRE displayed a lower and similar value relative to CT1 and CT2, respectively. For P2-EMRE, ATP-linked OCR did not differ from CT1 and CT2. In case of P3-MICU1, no difference in ATP-linked OCR with the other cell lines was observed. Explorative plotting of routine ECAR as a function of routine OCR (**Fig. 5H**) suggested that P3-MICU1 cells displayed a higher metabolic activity (*i*.*e*. an increased OCR and ECAR). Taken together, these results suggest that OXPHOS and glycolytic activity are not affected in P1-EMRE and P2-EMRE cells.

## Discussion

This study presents the integrated analysis of two novel *SMDT1* variants at the genetic, clinical and cellular level. The *SMDT1* gene encodes EMRE, which is a key regulating protein of the MCU complex and thereby of mitochondrial Ca^2+^ uptake. Both *SMDT1* variants induced absence of EMRE protein but not of MICU1, associated with formation of an MCU subcomplex of lower MW and impaired mitochondrial Ca^2+^ uptake. Importantly, complementation of the *SMDT1*-patient cells with the wild type *SMDT1* gene induced formation of a “normal” MCU complex and reversed the aberrant mitochondrial Ca^2+^ uptake phenotype, providing further evidence that the *SMDT1* variants are responsible for the observed cellular phenotype.

### SMDT1 variants induce absence of the EMRE protein and formation of an MCU subcomplex

The two patients presented in this study harboured a frameshift variant (P1-EMRE) or a missense variant (P2-EMRE) in the *SMDT1* gene. Western blot (WB) and immunofluorescence (IF) analysis provided evidence that MICU1 was present whereas EMRE was absent in *SMDT1*-patient cells. In P1-EMRE cells this absence was expected, since the variant was a frameshift that introduced a premature stop codon. In case of P2-EMRE cells, the detected EMRE missense variant (p.(Pro60Leu)) changes a key amino acid likely mediating the interaction between EMRE and MCU (69)(**Yamamoto et al., 2016**). Loss of this interaction causes rapid EMRE degradation (70, 71)(**König et al., 2016; Tsai et al., 2017**), compatible with the EMRE protein being absent in P2-EMRE cells (**Fig. 1A**). In case of the *MICU1* variant (a premature stop codon), MICU1 was absent but mitochondrial EMRE was detected (likely at reduced levels) by an IF-optimized antibody (**Fig. 2**). Mitochondrial EMRE was not detected in P3-MICU1 cells using SDS-PAGE analysis and a WB-optimized antibody (**Fig. 1A**). Likely, this discrepancy can be explained by the difference between the two antibodies. The IF-optimized antibody was raised against a larger immunogen sequence than the WB-optimized antibody (*i*.*e*. ISKNFAALLEEHDIFVPEDDDDDD *vs*. EEHDIFVPEDDD) and as such may show a higher affinity for the EMRE protein or recognize a slightly different epitope. Indeed, P3-MICU1 cells still displayed mitochondrial Ca^2+^ uptake and it is known that EMRE is essential for MCU-mediated Ca^2+^ uptake (39)(**Wang et al., 2019**). This suggests that *in situ*, absence of MICU1 induces formation of a destabilized (EMRE-containing) MCU complex, which is still able to mediate mitochondrial Ca^2+^ uptake (**Fig. 3B**). This agrees with previous evidence demonstrating that EMRE levels are lower upon MICU1 knockout (37, 50)(**Liu et al., 2016; Garg et al., 2021**). However, our results argue against a mechanism in which absence of MICU1 increased EMRE levels in *MICU1* patients (60)(**Bhosale et al., 2017**). Although differences in VDAC protein expression level between cell lines are obvious (**Fig. 1**), these cannot explain the impaired mitochondrial calcium uptake, since complementation with the wildtype *SMDT1* gene alleviated this phenotype (**Fig. 4**). Taken together, the provided evidence suggests that the studied mutations induce complete (P1-EMRE, P2-EMRE) or partial (P3-MICU1) absence of EMRE protein, leading to MCU destabilization. This conclusion is supported by previous animal and cell studies demonstrating that loss of EMRE or MICU1 induces formation of an MCU subcomplex (27, 28, 38, 43, 70)(**Sancak et al., 2013; Plovanich et al., 2013; König et al., 2016; Liu et al., 2020; Payne et al., 2020**).

### SMDT1 variants impair mitochondrial Ca^2+^ uptake during hormone stimulation but do not alter key parameters of mitochondrial function and morphology

Functional analysis of BK-stimulated PHSFs demonstrated that mitochondrial Ca^2+^ uptake was impaired in both *SMDT1*-patient cell lines. Reintroduction of the wild type *SMDT1* gene in P1-EMRE cells mitigated MCU subcomplex formation (**Fig. 4C**) and aberrant mitochondrial Ca^2+^ uptake (**Fig. 4D**), supporting the above conclusion that the absence of EMRE protein is causally linked to MCU-subcomplex formation and impairment of MCU-mediated mitochondrial Ca^2+^ uptake. The latter was not due to ΔΨ depolarization, compatible with other EMRE deficiency models (28, 42, 67)(**Sancak et al., 2013, Gottschalk et al., 2019; Tufi et al., 2019**). *SMDT1*-patient cells displayed normal enzymatic OXPHOS/CS activity (**Table 1**), normal routine OCR (**Fig. 5E**), normal ATP-linked OCR (**Fig. 5F**), and normal routine ECAR (**Fig. 5G**). These results are compatible with other studies demonstrating that: (**1**) *Smdt1*^*-/-*^ flies or mice exhibit normal routine mitochondrial respiration (42, 43)(**Tufi et al., 2019; Liu et al., 2020**), (**2**) tissue-specific *Mcu* ^*-/-*^ mice display normal routine mitochondrial respiration in cardiac and skeletal muscle tissue (47, 48)(**Kwong et al., 2015; Kwong et al., 2018**), and (**3**) routine OCR is normal in mouse embryonic fibroblasts or skeletal muscle mitochondria from whole-body *Mcu*^*-/-*^ mice (44)(**Pan et al., 2013**). Although P3-MICU1 cells displayed the previously reported fragmented mitochondrial phenotype (51, 67)(**Logan et al., 2014; Gottschalk et al., 2019**), no alterations in mitochondrial morphology were observed in P1-EMRE and P2-EMRE cells (**Fig. 5B-C-D**). Collectively, these findings demonstrate that hormone-stimulated, MCU-mediated, mitochondrial Ca^2+^ uptake is impaired in *SMDT1*-patient cells, but that key readouts of mitochondrial metabolic function are not altered. Regarding P3-MICU1, we observed that a large fraction of P3-MICU1 cells (71%) displayed increased mitochondrial rhod-2 fluorescence prior to hormone stimulation. Previous work demonstrated that MCU-mediated mitochondrial Ca^2+^ uptake stimulates mitochondrial OCR (72, 73)(**Llorente-Folch et al., 2013; Rueda et al., 2014**). Moreover, *Micu1*^*-/-*^ mice displayed increased serum lactate levels (50)(**Liu et al., 2016**), a phenomenon typically associated with increased glycolytic activity (68, 74)(**Divakaruni et al., 2014; Liemburg-Apers et al., 2015**). In this way, increased mitochondrial Ca^2+^ uptake might be responsible for the increased OCR and ECAR in non-stimulated P3-MICU1 cells, compatible with a higher metabolic activity (**Fig. 5H**).

### Potential pathomechanism in patients harbouring SMDT1 variants

Although the clinical phenotype of P1-EMRE and P2-EMRE only partially overlapped, both patients display skeletal muscle breakdown, as evidenced by elevated CK levels. In this context, P1-EMRE differs from P2-EMRE in that CK levels were also elevated during symptom-free intervals. Especially, P2-EMRE suffers from severe proximal muscle weakness displaying a “limb-girdle” distribution, which is in analogy to the alleged LGMD reported for a previously reported potential EMRE-variant (54)(**Monies et al., 2017**). In mice, MCU-mediated mitochondrial Ca^2+^ uptake counteracts the pathological loss of muscle mass and is involved in muscle trophism (78, 79)(**Mammucari et al., 2015; Gherardi et al., 2019**). In aged human subjects, training apparently improved muscle function, accompanied by an increase in MCU protein level (80)(**Zampieri et al., 2016**). In this sense, a lack of proper MCU complex function might be responsible for the enhanced skeletal muscle breakdown observed in P1-EMRE and P2-EMRE patients. Mitochondria are essential for proper skeletal muscle function as they supply the necessary ATP for actomyosin contraction and SERCA-mediated Ca^2+^ re-uptake into the sarcoplasmic reticulum (SR; (81)**de Groof et al., 2002**). The clinical phenotype of P1-EMRE is relatively mild but deteriorated considerably under stress conditions (*i*.*e*. during infection). Also, P2-EMRE reported an infection as the trigger for an observed deterioration in muscle strength. We therefore propose that deficient Ca^2+^-stimulated mitochondrial ATP generation might play a role in the stress-induced deterioration of the clinical phenotype in P1-EMRE and P2-EMRE. Although the *SMDT1* patients did not display classical signs of mitochondrial disease (*i*.*e*. lactic acidosis or enzymatic OXPHOS deficiencies), their stress sensitivity compares well to Leigh Syndrome patients with OXPHOS mutations and a reduced mitochondrial ATP production capacity, where (mild) intercurrent illnesses trigger disease exacerbation (82)(**Smeitink et al., 2004**). Several studies reported a reduced exercise capacity and impaired stress response in *Mcu*^*-/-*^ mice (44, 46-48)(**Pan et al., 2013; Luongo et al., 2015; Kwong et al., 2015; Kwong et al., 2018**). Although born at a much lower rate than predicted form Mendelian genetics, *Smdt1*^*-/-*^ mice behaved normally and did not differ from their wildtype littermates in their capability to perform maximal work or respond to acute stress (43)(**Liu et al., 2020**). These data are in line with the phenotype of P1-EMRE and P2-EMRE cells at the level of OXPHOS activity. Murphy and co-workers provide evidence that loss of EMRE induces compensatory adaptive pathways (43))(**Liu et al., 2020**), compatible with the observation that embryonic lethality depended on genetic background in *Mcu*^*-/-*^ and *Smdt1*^*-/-*^ mice (43-46)(**Luongo et al., 2015, Liu et al., 2020**). It remains to be determined whether the differences in the clinical phenotypes observed between P1-EMRE and P2-EMRE are linked to differences in genetic background and/or age. Future identification of additional *SMDT1*-patients will help to more precisely delineate the clinical spectrum associated with this gene defect.

## Materials and Methods

### Culture of primary human skin fibroblasts

Primary human skin fibroblasts (PHSFs) were obtained from skin biopsies. Informed consent for diagnostic and research studies was obtained for all subjects in accordance with the Declaration of Helsinki following the regulations of the local medical ethics committee. Cell lines used in this study were (**Table 1**): (**1**) control cell lines CT1 and CT2 obtained from skin biopsies of two healthy volunteers, (**2**) P1-EMRE and P2-EMRE cell lines obtained from skin biopsies of two patients with genetic variants in the EMRE-encoding *SMDT1* (single-pass membrane protein with aspartate rich tail 1) gene and, (**3**) a P3-MICU1 cell line obtained from a skin biopsy of a patient with an established pathogenic variant in the *MICU1* (mitochondrial calcium uptake 1) gene. PHSFs were cultured in medium 199 (M199; #22340-020; **Gibco Thermo Fisher Scientific, Waltham, MA, USA**) supplemented with 10% (v/v) Fetal Bovine Serum (FBS; #10270-106; lot #42Q2450K; **Gibco**) and 100 IU/mL penicillin/streptomycin (#15140122, **Gibco**) in a humidified atmosphere consisting of 95% air and 5% CO_2_ at 37°C. All experiments with PHSFs were performed at passage numbers between 8 and 28. Cells were regularly tested and found negative for mycoplasm using the MycoAlert mycoplasma detection kit (**Lonza**).

### Whole exome sequencing

Whole exome sequencing (WES) and data analysis were performed as described previously (83, 84)(**Neveling, 2013; Wortmann et al., 2015**). Briefly, exome enrichment was performed using the SureSelect Human All Exon 50 Mb Kit V5 (**Agilent Technologies, Santa Clara, CA, USA**). Sequencing was done on a HiSeq4000 (**Illumina, San Diego, CA, USA**) with a minimum median coverage of ×80. Read alignment to the human reference genome (GRCh37/hg19) and variant calling was performed at BGI (**Copenhagen, Denmark**) using BWA (Burroughs-Wheeler Aligner) and GATK (The Genome Analysis Toolkit) software, respectively. Variant annotation was performed using a custom designed in-house annotation. Intronic variants (except for splice sites), synonymous changes, and common variants were filtered and excluded from the initial datasets. Patient data were first analysed using a custom-made virtual gene panel consisting of known disease genes (as described in OMIM: https://omim.org) associated with muscle, mitochondrial, metabolic, and movement disorder disease (patient 1 = P1-EMRE) or muscle disease (patient 2 = P2-EMRE). As no disease-causing variants were detected, the entire exome was investigated for rare, protein damaging variants. This was done by comparison with gnomAD (The Genome Aggregation Database; gnomad.broadinstitute.org), dbSNPv132 and our in-house variant database with the MAF (minor allele frequency) depending on the mode of inheritance.

### Generation of complemented cell lines

Lentiviral complementation was performed as described in detail previously (85, 86)(**Baertling et al., 2017a; Baertling et al., 2017b**). In brief, wild type *SMDT1* (NCBI reference NM_033318.4) or green fluorescent protein (*GFP*; “Emerald GFP”) were cloned into a pLenti6.2/V5 destination vector carrying the blasticidin resistance gene, using Gateway technology (**Invitrogen Thermo Fisher Scientific, Waltham, MA, USA**), creating a *SMDT1* or *GFP* open reading frame with a C-terminal V5-tag. Lentiviral particles were produced by transfecting HEK293T cells (#632180; **Clontech, Westburg, Leusden, The Netherlands**) according to the manufacturer’s protocol (**Invitrogen**). Virus particles were harvested 72 h after transfection and added to control (CT1) and patient (P1-EMRE) PHSFs. After 24 h, the virus-containing medium was replaced by a virus-free medium. After 48 h, blasticidin (2 μg/ml; # ant-bl-1; **InvivoGen, San Diego, USA**) was added to the medium to select for stably transduced SMDT1-V5 or GFP-V5 cells. Control- and patient-derived cells (CT1+GFP-V5, CT1+SMDT1-V5, P1-EMRE+GFP-V5, P1-EMRE+SMDT1-V5) were cultured in M199 medium supplemented with 10% (v/v) Fetal Bovine Serum (FBS; #10270-106; lot #42Q2450K; **Gibco**) and 100 IU/mL penicillin/streptomycin (#15140122, **Gibco**) in a humidified atmosphere consisting of 95% air and 5% CO_2_ at 37°C. Prior to analysis, the cells were cultured in the presence of 2 μg/ml blasticidin for 14 days. CT1 and P1-EMRE cells were transduced at passage 17 and 8, respectively. Transduced PHSFs were used between passage 17+8 and 17+15 (CT1+GFP-V5 and CT1+SMDT1-V5) and 8+8 and 8+21 (P1+GFP-V5 and P1+SMDT1-V5).

### Preparation of mitochondria-enriched fractions

PHSF pellets (∼10×10^6^ cells) were resuspended in 1250 μl Tris-HCl buffer (10 mM, pH 7.6). This cell suspension was homogenized using a Potter-Elvehjem homogenizer (8-10 strokes at 1800 rpm on ice) and then made isotonic by addition of 250 μl sucrose (1.5 M; final concentration 250 mM). Next, two centrifugation steps were used to remove cell debris (10 min, 600 g, 4°C) and to isolate a mitochondria-enriched fraction from the supernatant (10 min, 14,000 g, 4°C). The resulting pellet was resuspended in Tris-HCl (10 mM; pH 7.6) and stored at −80°C until further analysis.

### Enzymology of oxidative phosphorylation enzymes

The enzyme activities of oxidative phosphorylation (OXPHOS) complexes I-V (CI-CV) and citrate synthase (CS) were determined in PHSF-derived mitochondria-enriched fractions as described previously (87)(**Rodenburg, 2011**). All activities were normalized to total mg protein or CS activity (**Table 1**).

### SDS-PAGE and Western blotting (WB)

Cells were harvested by trypsinization, washed with cold PBS, centrifuged (5 min, 1000 g, 4°C) and resuspended in 250 μL MSE buffer (225 mM mannitol, 75 mM D-sucrose and 1 mM Na-EDTA, pH 7.4) supplemented with 1x protease inhibitor cocktail (#05892791001; **Roche Diagnostics Merck**). Cells were exposed to three cycles of cold (liquid nitrogen) and heat shock (37°C) and homogenized with a micro pestle. Cell debris was pelleted by centrifugation (15 min, 600 g, 4°C). The supernatant was centrifuged at high speed in order to pellet mitochondria (15 min, 10,000 g, 4°C). To serve as cytosolic fraction, 200 μL of the supernatant was saved. The mitochondrial pellet was dissolved in 40 μL PBS containing 2% (w/v) β-lauryl maltoside and incubated on ice for 10 minutes. Protein concentrations were determined using Protein Assay Dye Reagent Concentrate (#500-0006; **Bio-Rad**). Spectrophotometric absorbance was measured at 595 nm in a Benchmark Plus plate reader (**Bio-Rad**). Samples were prepared for loading by mixing 20-40 μg of mitochondrial and cytosolic fractions with 5 x Sample Buffer (in Milli Q) containing: 0.8 M DDT, 10% (w/v) SDS, 250 mM Tris-HCl pH 6.8, 60% (v/v) glycerol, and 0.03% (w/v) bromophenol blue (#1610404; **Bio-Rad**). Next, proteins were separated on a 4-15% Mini-PROTEAN TGX Stain-Free SDS-PAGE gel (#456-8084; **Bio-Rad**) and transferred to a 0.2 μm PVDF membrane (#ISEQ00010; **Merck**) using a standard ice-cold wet blotting system (**Bio-Rad**). The membranes were blocked with Intercept Blocking Buffer (#927-70001; **Li-cor**) supplemented with 0.1% (v/v) Tween-20 for 1 h at RT. Subsequently, the blots were incubated with one of the following primary antibodies in Intercept Blocking Buffer supplemented with 0.1% (v/v) Tween-20 overnight at 4°C: an WB-optimized EMRE antibody (anti-EMRE; #A300-BL19208, 1:250; **Bethyl Laboratories, Montgomery, TX, USA**), anti-MICU1/CALC/CBARA1 (#LN2014887, 1:2000; **Labned, Amstelveen, The Netherlands**), anti-MCU/C1orf42/CCDC109A (#26312-1-AP, 1:2000; **Proteintech, Rosemont, IL, USA**), anti-VDAC1/Porin (#MABN504, 1:1000; **Merck**), anti-β-Actin/ACTB (#A5441, 1:100,000; **Merck**), anti-V5 (#R960-25; 1:5000; **Thermo Fischer**). Next, membranes were incubated with secondary antibodies goat anti-rabbit IRDye800 (#926-32211; **Westburg, Leusden, The Netherlands**) and goat anti-mouse IRDye680 (#926-68070; **Westburg**) diluted (both 1:10,000) in Intercept Blocking Buffer supplemented with 0.1% (v/v) Tween-20 for 1 h at room temperature. The blot was scanned using an Odyssey CLx scanner (**Li-cor**). In case of the V5 tag, mitochondria-enriched fractions were processed by adding 0.25% (v/v) of SDS-PAGE sample buffer (#PCG3009; **Sigma-Aldrich/Merck**) to 20 μg of mitochondrial protein and run with Tris-MOPS running buffer (#PCG3003; **Sigma-Aldrich/Merck**) on a 4-20% precast Trupage gel (#PCG2012-10EA; **Sigma-Aldrich/Merck**). For Western blotting, proteins were transferred to a 0.2 μm PVDF membrane (#ISEQ85R; **Sigma-Aldrich/Merck**) by electroblotting the separated proteins for 1 h at 100 V. Antibody incubation and visualization of chemiluminescence signal was performed as described for native gel electrophoresis below.

### BN-PAGE and Western blotting (WB)

Mitochondria-enriched fractions (30 μg total protein) were suspended in a solution containing 50 mM Bis/Tris, 50 mM NaCl, 10% (v/v) glycerol and 0.001% (w/v) Ponceau S (#114275; **Merck**); pH was adjusted to 7.2 with HCl. To solubilize mitochondrial proteins, digitonin (#19551; **Serva**) was added in a 10:1 (w/w) digitonin-to-protein ratio (20 min on ice). Next, this solution was centrifuged (30 min; 20,000 g; 4°C) and Serva Blue G (#35050; **Serva**) was added to the supernatant in a 1:5 (w/w) ratio relative to digitonin. Blue native PAGE (BN-PAGE) was performed using 3-12% precast NativePAGE Bis-Tris gels (#BN1001BOX; **Invitrogen**). For western blotting, proteins were transferred to a 0.45 μm PVDF membrane (#IPVH85R; **Sigma-Aldrich/Merck, Zwijndrecht, The Netherlands**) using an electroblotting device (**Bio-Rad**; 100 V, 1 h). Membranes were blocked using 2% (w/v) non-fat milk (NFM) in Tris-buffered saline with 0.1% (v/v) Triton (TBS-Triton). Subsequently, blots were incubated with primary antibodies in 3% (w/v) BSA/TBS-Triton for 2 h. Antibodies included: anti-MCU (#26312-1-AP; **Proteintech Europe, Manchester, UK**) or anti-CII (SDHA; #Ab14715; **Abcam, Cambridge, UK**). Next, the blots were incubated with secondary antibodies: horseradish peroxidase-conjugated goat anti-mouse (#P0047; **Dako Products Agilent, Santa Clara, CA, USA**) or goat anti-rabbit (#A00160; **Genscript Biotech, Piscataway, NJ, USA**) in 2% (w/v) NFM/TBS-Triton for 1 h. Chemiluminescence signals were visualized using the enhanced chemiluminescence kit (ECL; #32106; **Thermo Fischer**) and a Chemidoc XRS+ scanner system (**Bio-Rad, Hercules, CA, USA**).

### Immunofluorescence microscopy (IF)

Fibroblasts were seeded in an 8-well glass-bottomed chamber (#155411; **Nunc Lab-Tek Thermo Scientific, Waltham, MA, USA**), three days prior to the experiment. To visualize mitochondria, cells were incubated with Mitotracker Red-FM (0.5 μM in PBS; #M22425; **Thermo Fischer**) for 30 min at 37°C. Next, the cells were washed with PBS, fixed with 4% (w/v) paraformaldehyde (20 min), permeabilised with 0.3% (v/v) Triton X-100 (15 min) and blocked in PBS supplemented with 1% (w/v) BSA (PBS-BSA) for 30 min at RT. To visualize EMRE protein, cells were incubated with an IF-optimized EMRE antibody (anti-SMDT1/C22orf32/EMRE; #HPA060340, 1:250; **Merck**), washed 3 times in PBS, and incubated with a secondary Goat-anti-Rabbit antibody conjugated to Alexa-488 (1:400 in PBS-BSA for 1 h at RT in the dark; #A-11008; **Thermo Fischer**). Then the cells were washed in PBS and imaged in PBS-BSA using a TCS SP5 confocal microscope (**Leica Microsystems, Mannheim, Germany**) equipped with an HCX PL-APO 63x N.A. 1.2 water immersion objective at 37°C. Alexa-488 was excited with an Argon laser (488 nm) and emission was detected between 500-550 nm. MitoTracker-Red was excited using a DPSS laser (561 nm) and emission was detected between 572-674 nm.

### Mitochondrial Ca^2+^ uptake

Two days prior to imaging, PHSFs were seeded at 35,000 cells/dish on FluoroDishes (FD35-100, **World Precision Instruments, Sarasota, FL, USA**). Next, the cells were co-incubated with 3 μM of the cytosolic Ca^2+^ reporter molecule fura-2 AM (#F1221; **Thermo Fischer**) and 5 μM of the mitochondrial matrix Ca^2+^ reporter molecule rhod-2 AM (#R1245MP; **Thermo Fischer**) in HEPES-Tris buffer (HT; 132 mM NaCl, 4.2 mM KCl, 1 mM CaCl_2_, 1 mM MgCl_2_, 5.5 mM D-glucose and 10 mM HEPES, pH 7.4) for 20 min at 37°C. After removal of the reporter molecules, the cells were incubated for 10 min at 37°C in HT buffer to allow for full de-esterification and equilibration of the internalized reporter molecules. Next, the HT buffer was refreshed and the cells were placed in a temperature-controlled (37°C) plate attached to the stage of an inverted microscope (Axiovert 200M; **Zeiss, Jena, Germany**) equipped with a 40x F Fluar 1.3 N.A. oil immersion objective, and allowed to equilibrate for 5 min. Fura-2 and rhod-2 were alternatingly excited at 380 and 540 nm, respectively, using a monochromator (Polychrome IV; **TILL Photonics, Gräfelfing, Germany**). Fura-2 fluorescence light was directed by a 430 DCLP dichroic mirror (**Omega Optical, Brattleboro, VT**) through a 510wb40 emission filter (**Omega**) on a CoolSNAP HQ monochrome charge coupled device (CCD) camera (**Roper Scientific, Vianen, The Netherlands**). Rhod-2 fluorescence light was directed by a 560DRLP dichroic mirror (**Omega**) through a 565ALP emission filter (**Omega**). The camera exposure time was set at 50 ms using an image acquisition interval of 4 s. Experiments with P2-EMRE cells were carried out using a microscopy system consisting of a Polychrome 5000 monochromator (**TILL photonics**), an AxioObserver Z1 inverted microscope (**Zeiss**) and a CoolSNAP HQ2 camera (**Roper Scientific**) using similar dichroic mirrors and emission filters (**Omega**). All hardware was controlled using Metafluor 6.0 software (**Universal Imaging Corporation, Downingtown, PA**). During time measurements, cells were stimulated with the hormone Bradykinin (BK; 1 μM; #B3259, **Merck**) to induce mitochondrial Ca^2+^ uptake (6, 55-57)(**Visch et al., 2004; Visch et al., 2006a; Visch et al., 2006b; Valsecchi et al., 2009**). Cells were considered positive for mitochondrial Ca^2+^ uptake (scored by two different researchers in a blinded manner) when meeting all of the following criteria (*e*.*g*. **Fig. 3A**): (**I**) Prior to BK addition, mitochondrial rhod-2 emission is absent. P3-MICU1 cells were exempt from this criterion, since a *MICU1* gene defect can induce an increase in basal mitochondrial Ca^2+^ level (50, 51, 60)(**Logan et al., 2014; Liu et al., 2016; Bhosale et al., 2017**), (**II**) Cells responded to BK, as demonstrated by an acute decrease in the 380 nm fura-2 signal and an acute increase in the nuclear rhod-2 signal, and (**III**) a mitochondria-located rhod-2 fluorescence signal was observed upon BK addition. While imaging, the image focus was manually adjusted to guarantee that no mitochondrial rhod-2 signals were missed during BK-stimulation.

### Mitochondrial TMRM accumulation and morphology analysis

Three days prior to imaging, cells were seeded at 12,000 cells/dish on fluorodishes (#FD35-100, **World Precision Instruments**). Cells were incubated with 15 nM tetramethyl rhodamine methyl ester (TMRM; #T668; **Invitrogen**) in culture medium for 25 min at 37°C and 5% CO_2_ in the dark. Then, the dish was mounted on the Axiovert 200M microscope system described above and imaged in the culture medium containing 15 nM TMRM. TMRM was excited at 540 nm and emission light was directed by a 560DRLP dichroic mirror through a 565ALP emission filter. For each dish, 25-30 fields of view (FOVs) were acquired using an exposure time of 50-100 ms.

### Cellular oxygen consumption and extracellular acidification measurements

Oxygen consumption rate (OCR) and extracellular acidification rate (ECAR) of PHSFs were analysed using a Seahorse XFe96 analyzer (**Agilent**). On the day of the assay, a Cell Culture Microplate (#101085-004; Agilent, Santa Clara, CA, USA) was coated with Cell-Tak® (22.4 μg/ml in 0.1 M NaHCO_3_; #734-1081; **BD Biosciences, San Jose, CA, USA**) at 37°C (not CO_2_ corrected) for at least 1 h. Prior to the assay, cells were seeded at a density of 15,000 cells per well (6 replicates for each cell line) in assay medium (DMEM supplemented with 1 mM pyruvate, 2 mM L-Glutamine and 11 mM D-Glucose; pH adjusted to 7.4 with NaOH) and incubated at 37°C (not CO_2_ corrected) for 1 h. OCR and ECAR were recorded in untreated cells using three cycles (each consisting of 3 min of mixing followed by 3 min of recording). The numerical data of the last cycle was used to quantify basal (routine) OCR (OCR_routine_) and basal (routine) ECAR. Leak respiration (OCR_leak_) was determined similarly after addition of the F_o_F_1_-ATPase inhibitor oligomycin A (1 μM; #75351; **Sigma**). The ATP-linked OCR was determined by calculating OCR_routine_-OCR_leak_. Wells displaying a zero OCR value were excluded from the analysis.

### Data analysis

Image processing and quantification was performed using Image Pro Plus 6.1 (**Media Cybernetics, Rockville, MD, USA**) and FIJI software (https://fiji.sc). Numerical data is presented as mean±SEM (standard error of the mean) unless stated otherwise. Statistical significance was assessed using a non-parametric (Mann-Whitney U) test using Origin Pro 2020b software (**OriginLab Corp., Northampton, MA, USA**). Statistical significance is indicated by asterisks as follows: *p<0.05, **p<0.01 and ***p<0.001. Linear contrast stretch of immunofluorescence images was performed as described previously (*e*.*g*. (61)**Koopman et al., 2008**).

## Data availability

This study includes no data deposited in external repositories. Data analyzed during the current study is available from the corresponding authors on reasonable request.

## Acknowledgements

We thank Jozef Hertecant (Department of Pediatrics, Tawam Hospital, United Arab Emirates) for clinical diagnosis and care of patient 3 (P3-MICU1). We thank Laszlo Groh (Dept. of Internal Medicine, Radboudumc) and Els van de Westerlo (Dept. of Biochemistry, Radboudumc) for their practical assistance with Seahorse and Western blot experiments. We would like to thank the colleagues of the mitochondrial diagnostic group (muscle lab, cell culture lab and DNA lab) of the Translational Metabolic Laboratory at the Radboudumc for excellent technical assistance. We would like to acknowledge the Genome Technology Centre at the Radboudumc and BGI Copenhagen for technical support of the exome sequencing. We are grateful to Khondrion B.V. (Nijmegen, The Netherlands) for the use of their microscopy hardware.

## Abbreviations

ΔΨ: trans-MIM mitochondrial membrane potential
[Ca^2+^]_c_: cytosolic free Ca^2+^ concentration
[Ca^2+^]_m_: mitochondrial free Ca^2+^ concentration
[Ca^2+^]_n_: nuclear free Ca^2+^ concentration
BK: bradykinin
CK: creatine kinase
ECAR: extracellular acidification rate
EMRE: essential MCU Regulator
FOV: field of view
IF: immunofluorescence
LGMD: limb-girdle muscular dystrophy
MCU: mitochondrial calcium uniporter
MICU1/2: mitochondrial calcium uptake protein 1/2
MIM: mitochondrial inner membrane
MOM: mitochondrial outer membrane
MW: molecular weight
OCR: oxygen consumption rate
OXPHOS: oxidative phosphorylation
PHSFs: primary human skin fibroblasts
SMDT1: single-pass membrane protein with aspartate rich tail 1
TCA: tricarboxylic acid cycle
WB: Western blot
WES: whole exome sequencing.

