## Supplementary material for "*SMDT1* variants impair EMRE-mediated mitochondrial calcium uptake in patients with muscle involvement"

### Supplementary Table 1: Filtering of whole exome sequencing data of P1-EMRE and P2-EMRE

| Patient | **P1-EMRE** | **P2-EMRE** |
|---|---|---|
| DNA-no. | DNA16-07061B | DNA15-10039B |
| Variants | 138252 | 120936 |
| Allele frequency <1%* | 11901 | 3486 |
| Exonic/splice site variants | 2495 | 842 |
| Non-synonymous variants | 1532 | 531 |
| Variants compatible with recessive inheritance | 431 | 87 |
| Homozygous variants | 9 | 26 |
| Variants previously assessed as (likely) benign | 8 | 22 |
| Software scores predicting pathogenicity# | 1 | 2 |
| | **1 class 4 variant in *SMDT1*** | **1 class 3 variant in *SMDT1***<br><br>1 class 3 variant in *PISD* |
| | mean coverage: 144-fold | mean coverage: 96-fold |

*Allele frequency data in GnomAD

#Variant analysis software used: PhyloP, CADD, SIFT, Grantham

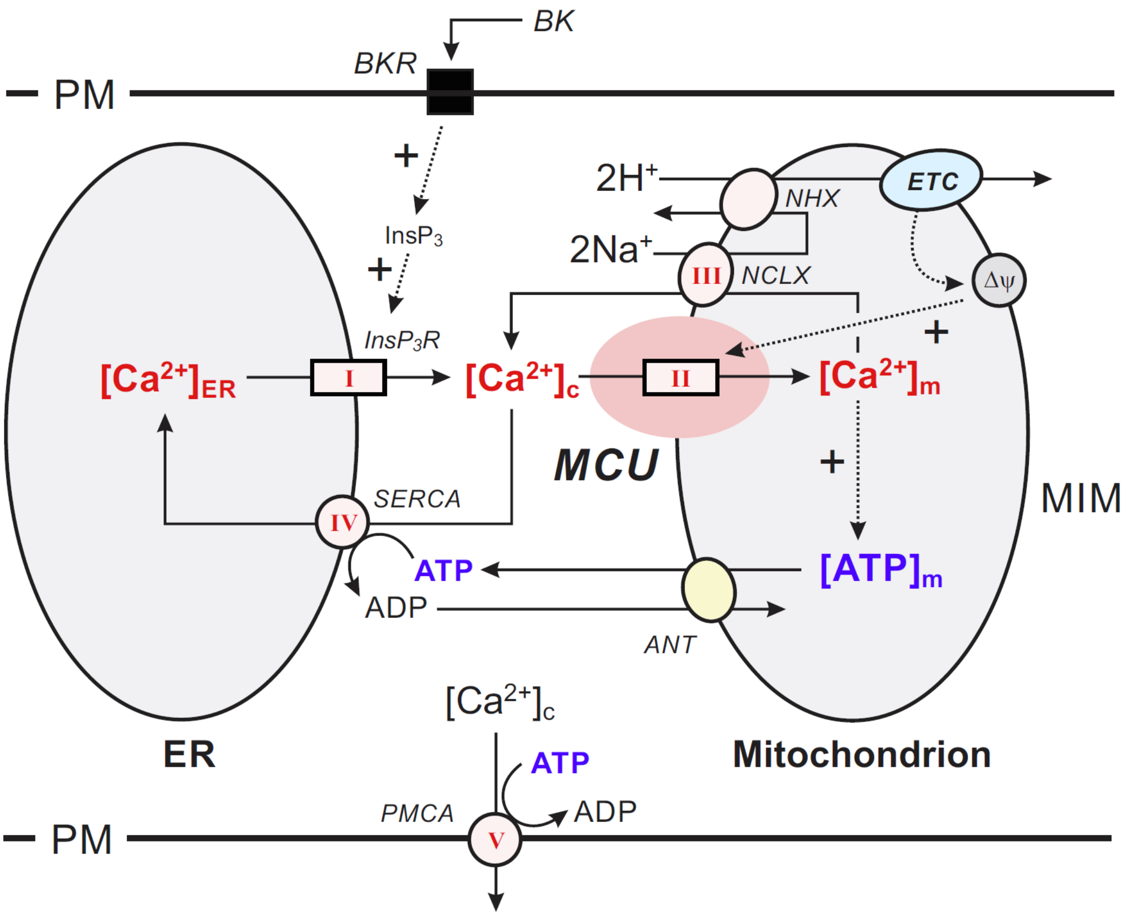


**Supplementary Figure 1: Calcium handling in hormone-stimulated PHSFs.** Stimulation of primary human skin fibroblasts (PHSFs) with the hormone Bradykinin (BK) releases calcium ($Ca^{2+}$) from the endoplasmic reticulum (ER) via stimulation (+) of the inositol 1,4,5-trisphosphate ($InsP_3$)-activated $InsP_3$-receptor ($InsP_3R$; ***pathway I***). The released $Ca^{2+}$ enters the mitochondrial matrix via MCU-mediated uptake (***pathway II***) and leaves this compartment again by the action of the mitochondrial sodium calcium exchanger (NCLX; ***pathway III***). The latter is coupled to the action of the mitochondrial sodium hydrogen exchanger (NHX) and the mitochondrial electron transport chain (ETC). The ETC also sustains the inside-negative electrical potential (Δψ) across the mitochondrial inner membrane (MIM), which stimulates (+) MCU-mediated mitochondrial $Ca^{2+}$ uptake. Within the mitochondrial matrix, $Ca^{2+}$ stimulates the generation of ATP, which is exported to the cytosol by the Adenine nucleotide translocator (ANT) to fuel SERCA-mediated ER $Ca^{2+}$ uptake (***pathway IV***). Alternatively, $Ca^{2+}$ can be actively removed from the cytosol by PMCA action (***pathway V***). **Abbreviations:** ADP, adenosine diphosphate; $[ATP]_m$, free ATP concentration in the mitochondrion; BKR, Bradykinin receptor; $[Ca^{2+}]_c$, free $Ca^{2+}$ concentration in the cytosol; $[Ca^{2+}]_{ER}$, free $Ca^{2+}$ concentration in the ER; $[Ca^{2+}]_m$, free $Ca^{2+}$ concentration in the mitochondrion; MCU, mitochondrial uniporter; PM, plasma membrane; SERCA, sarco/endoplasmic reticulum $Ca^{2+}$-ATPase; PMCA, plasma membrane $Ca^{2+}$-ATPase.

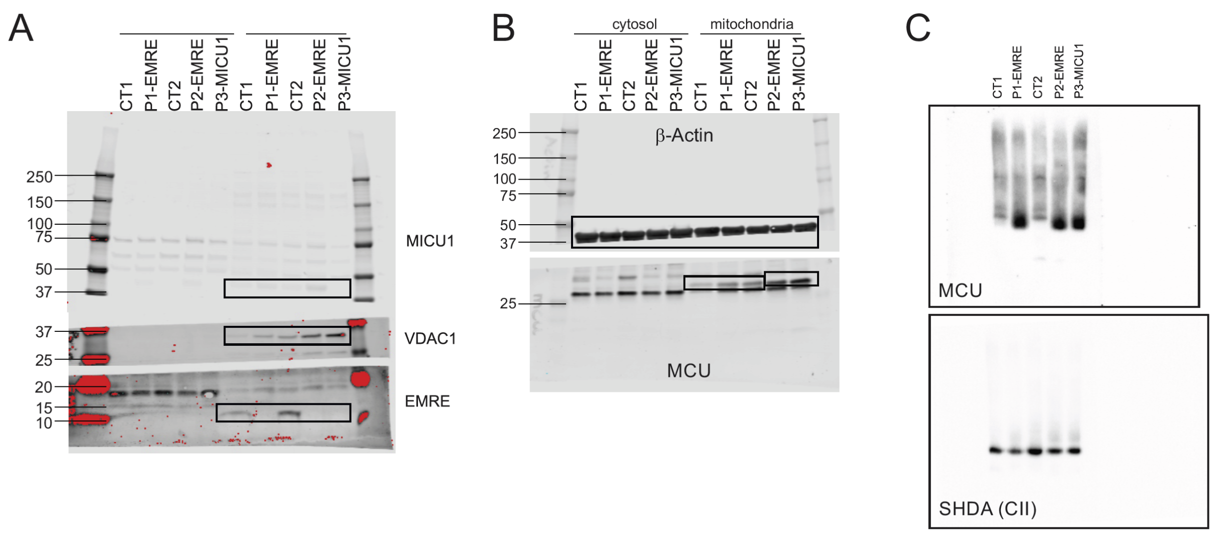


**Supplementary Figure 2: Original blots used for Figure 1.** Panel A and B were used to create **Fig. 1A.** Panel C was used to create **Fig. 1B**. numerals indicate molecular weight (MW) in kDa. Saturated pixels (markers) are depicted in red.

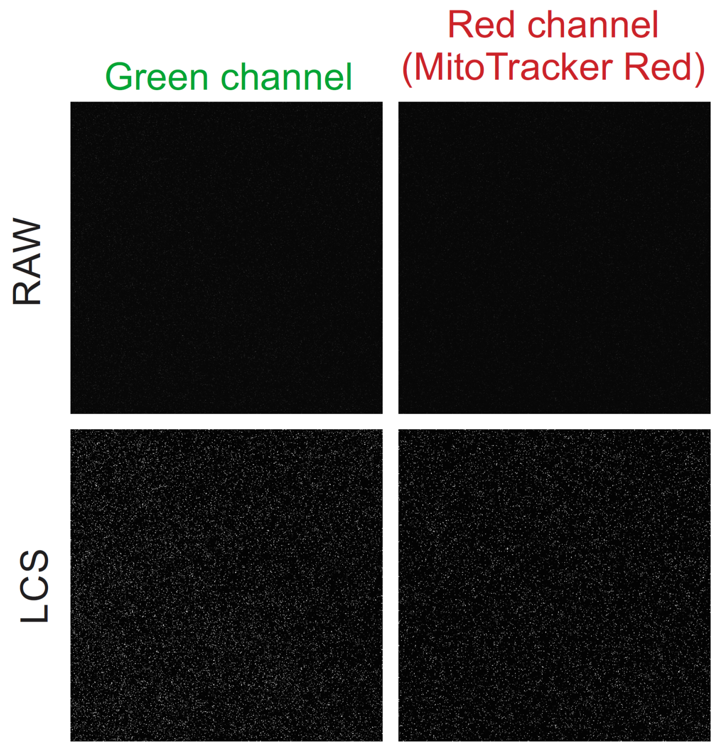


**Supplementary Figure 3A: Secondary antibody control for immunofluorescence.** Confocal microscopy images of CT2 cells. Cells were stained with anti-rabbit-Alexa-488 only in order to rule out the possibility of non-specific binding of the secondary antibody. Raw images are shown in the upper panel. Images in the lower panel were processed for visualization purposes by subsequent application of a linear contrast stretch (LCS) operation, a median filter (3x3; single pass) and a second LCS operation.

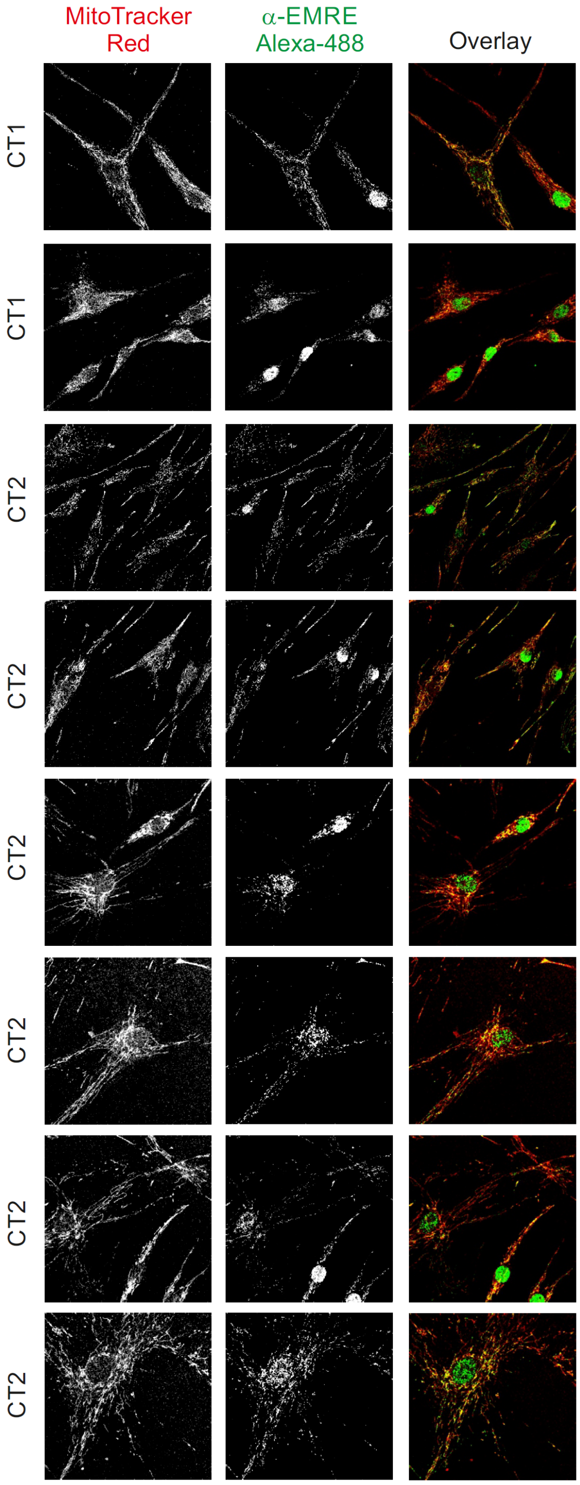


**Supplementary Figure 3B:** Additional confocal microscopy images of CT1 and CT2 cells co-stained with the mitochondrial marker MitoTracker Red (red) and anti-EMRE/Alexa-488 antibodies (green). Images were processed for visualization purposes by subsequent application of a linear contrast stretch (LCS) operation, a median filter (3x3; single pass) and a second LCS operation.

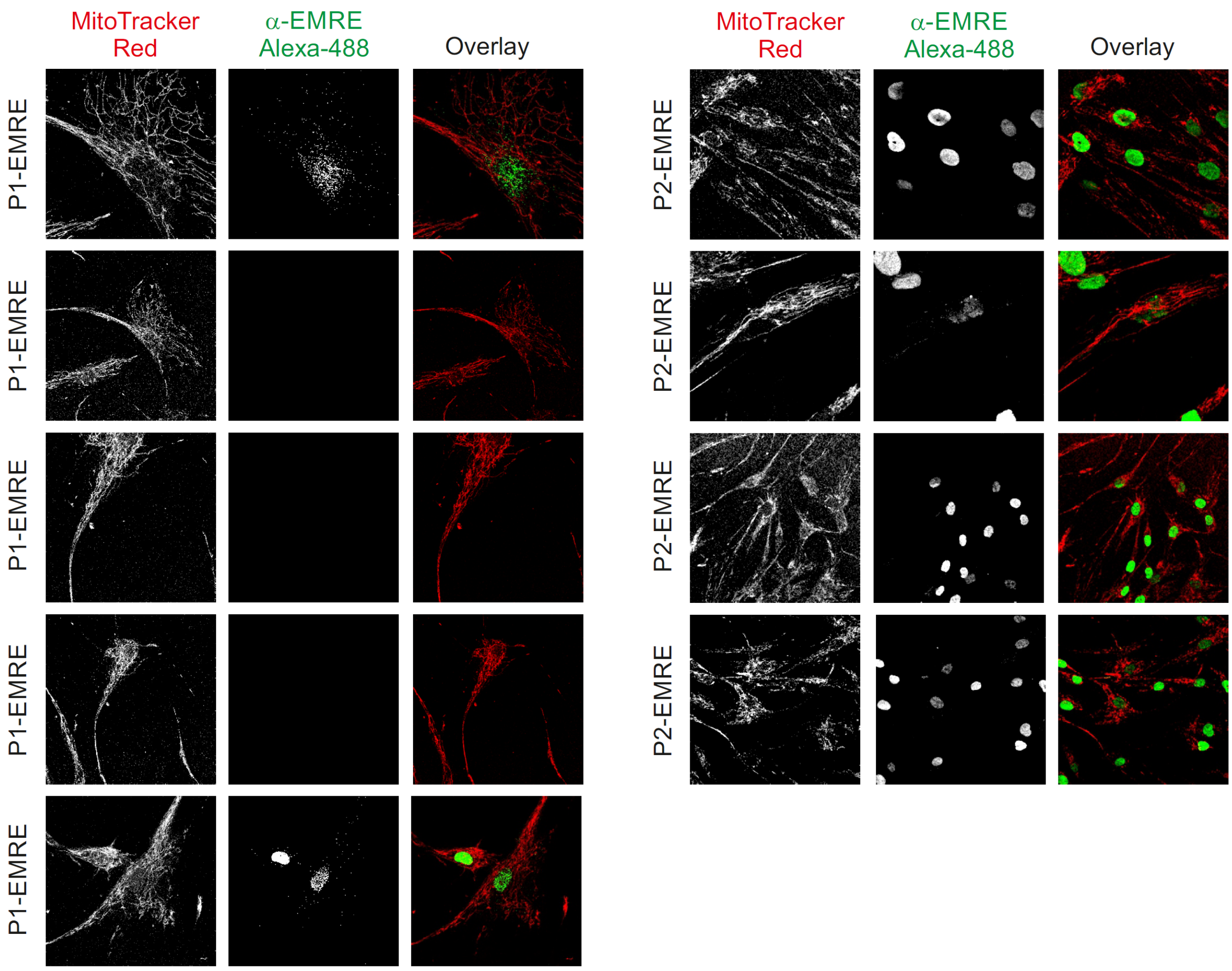


**Supplementary Figure 3C:** Additional confocal microscopy images of P1-EMRE and P2-EMRE cells co-stained with the mitochondrial marker MitoTracker Red (red) and anti-EMRE/Alexa-488 antibodies (green). Images were processed for visualization purposes by subsequent application of a linear contrast stretch (LCS) operation, a median filter (3x3; single pass) and a second LCS operation.

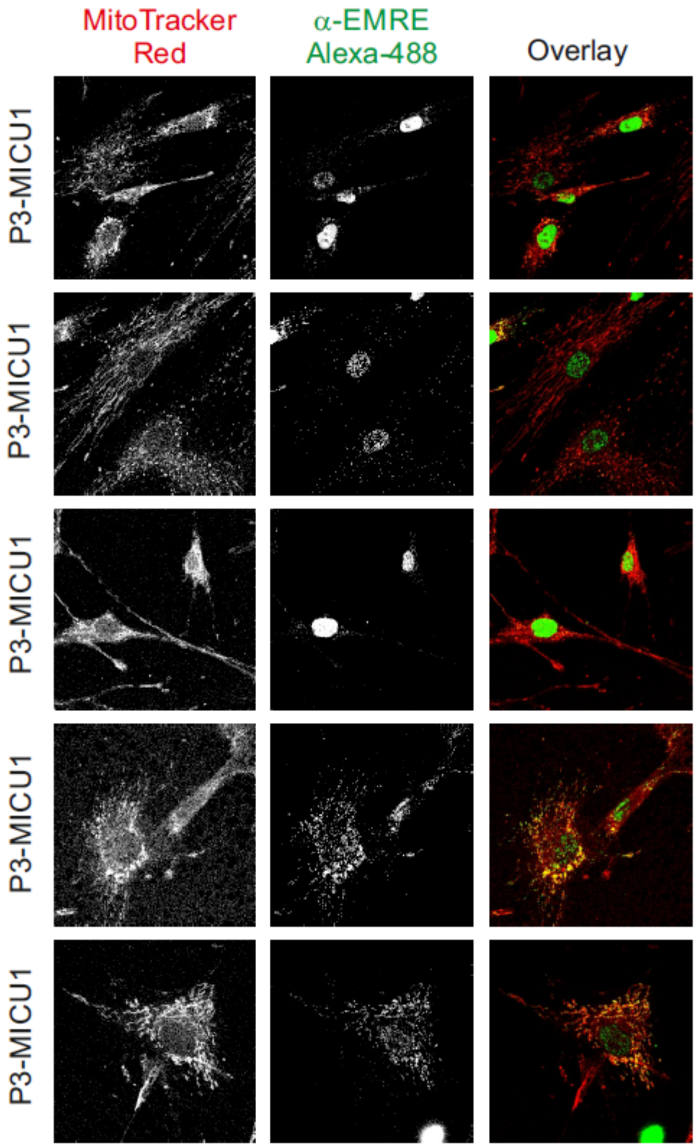


**Supplementary Figure 3D: EMRE antibody staining in P3-MICU1.** Additional confocal microscopy images of patient-derived PHSFs (P3-MICU1) co-stained with the mitochondrial marker MitoTracker Red (red) and anti-EMRE/Alexa-488 antibodies (green). Images were processed for visualization purposes by subsequent application of a linear contrast stretch (LCS) operation, a median filter (3x3; single pass) and a second LCS operation.

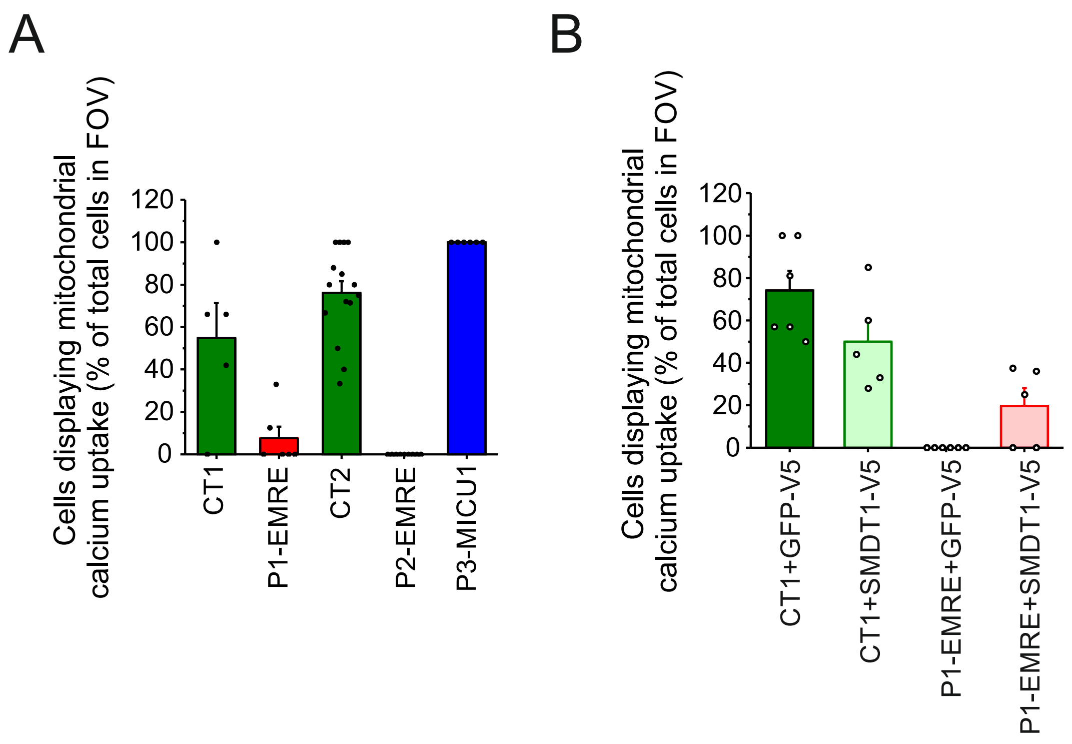


**Supplementary Figure 4: Second blinded analysis of the mitochondrial calcium uptake data.** (**A**) Scoring of BK-stimulated mitochondrial $Ca^{2+}$ uptake data in **Fig. 3B** by a second researcher. Each symbol represents a field of view (FOV). For blinding purposes, here, the number of FOVs counted was 15 for CT2. (**B**) Same as panel A, but now for the data used in **Fig. 4D**.

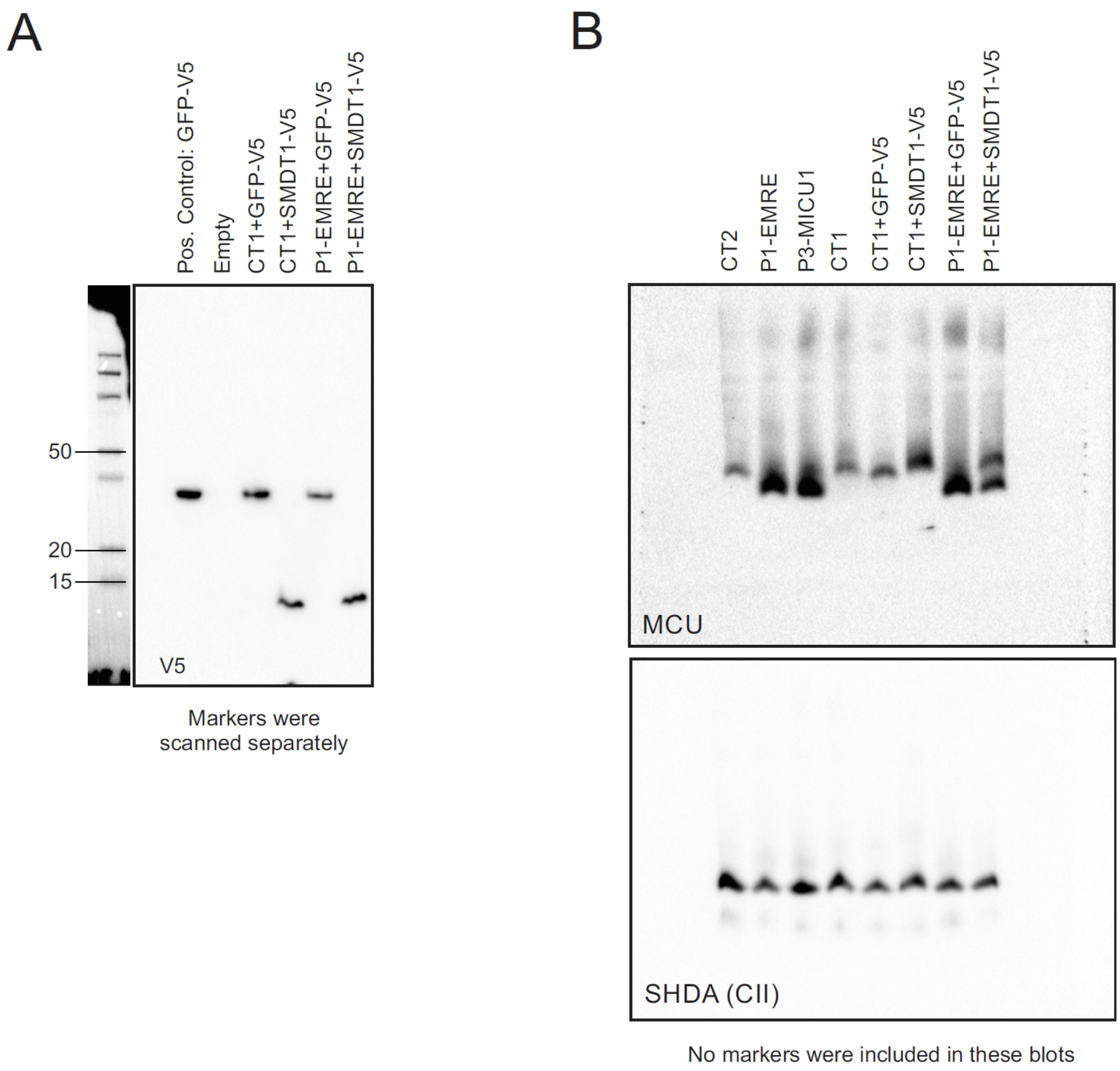


**Supplementary Figure 5: Original blots used for Figure 4.** (**A**) Data used for **Fig. 4A**. Numerals indicate molecular weight (MW) in kDa. (**B**) Data used for **Fig. 4C** (Lanes 5 to 8). For comparison some samples also used in **Fig. 1A** (CT1, CT2, P1-EMRE and P3-MICU1) were included in the analysis (lanes 1 to 4).

### MOVIES

**Supplementary movie 1: mitochondrial $Ca^{2+}$ uptake in CT1 cells.** Contrast-optimized (using ImagePro software) representative movie (created using Fiji) of CT1 cells loaded with fura-2 (3 µM; left) and rhod-2 (5 µM; right). The probes were alternately excited with 380 nm (fura-2) and 540 nm light (rhod-2) every 4 seconds. Bradykinin (1 µM) was added after ~25 cycles (104 sec). The focus was manually adjusted during the measurement to make sure that no out of focus mitochondrial rhod-2 signals were missed. The analysis strategy is explained in detail in the caption of **Figure 3A**.

**Supplementary movie 2: mitochondrial $Ca^{2+}$ uptake in P1-EMRE cells.** Contrast-optimized (using ImagePro software) representative movie (created using Fiji) of P1-EMRE cells loaded with fura-2 (3 µM; left) and rhod-2 (5 µM; right). The probes were alternately excited with 380 nm (fura-2) and 540 nm light (rhod-2) every 4 seconds. Bradykinin (1 µM) was added after ~25 cycles (104 sec). The focus was manually adjusted during the measurement to make sure that no out of focus mitochondrial rhod-2 signals were missed. The analysis strategy is explained in detail in the caption of **Figure 3A**.
